# Longitudinal modality prediction learns gene regulatory patterns: insights from a single-cell competition

**DOI:** 10.64898/2026.02.24.707614

**Authors:** Christopher Lance, Vladimir A. Shitov, Hongzhi Wen, Yuge Ji, Peter Holderrieth, Yuesong Wu, Renming Liu, Robrecht Cannoodt, Wenzhuo Tang, Kai Waldrant, Benjamin DeMeo, Mauricio Cortes, Daniel Kotlarz, Jiliang Tang, Yuying Xie, Fabian J. Theis, Daniel B. Burkhardt, Malte D. Luecken

**Affiliations:** Department of Computational Health, Institute of Translational Genomics, Helmholtz Munich, Munich, Germany; Dr. von Hauner Children’s Hospital, Department of Pediatrics, University Hospital, Ludwig-Maximilians-Universität Munich, Munich, Germany; Department of Computational Health, Institute of Computational Biology, Helmholtz Munich, Munich, Germany; Comprehensive Pneumology Center (CPC) with the CPC-M bioArchive / Institute of Lung Health and Immunity (LHI), Helmholtz Munich; Member of the German Center for Lung Research (DZL), Munich, Germany; Department of Computer Science and Engineering, Michigan State University, East Lansing, Michigan, United States of America; School of Life Sciences, Technical University of Munich, Weihenstephan, Germany; Cellarity, Inc., United States of America; Department of Statistics and Probability, Michigan State University, East Lansing, Michigan, United States of America; Data Intuitive, Belgium; Data Mining and Modelling for Biomedicine group, VIB Center for Inflammation Research, Ghent, Belgium; Department of Applied Mathematics, Computer Science, and Statistics, Ghent University, Ghent, Belgium; TUM School of Computation, Information & Technology, Technical University of Munich, Garching, Germany; TUM School of Life Sciences Weihenstephan, Technical University of Munich, Freising, Germany

## Abstract

Simultaneous measurement of chromatin, transcriptomic, and proteomic features in single cells opens new avenues for modeling interactions between molecular layers during dynamic biological processes. Predicting one modality from another - such as inferring gene expression from chromatin profiles or protein abundance from RNA - has the potential to reveal regulatory relationships and enhance downstream analyses. However, conventional approaches for predicting gene regulation have largely failed and method development in modality prediction for regulatory inference has been limited.

To explore effective modeling strategies and stimulate innovation, we generated a purpose-built longitudinal multimodal benchmarking dataset that captures early hematopoietic differentiation and organized the largest single-cell data competition to date, receiving over 27,000 submissions from 1,602 competitors worldwide. In our extensive analysis of the competition results, we demonstrated that top-performing approaches outperform state-of-the-art methods, and uncover how best-performing models captured biologically meaningful regulatory relationships between modalities. With ablation studies of the winning models, we identified feature-engineering strategies, model architectures and cross-validation schemes that are crucial for outstanding performance, and provide simplified, reproducible, light-weight code for state-of-the-art models. Together, the benchmark and analyses serve as an evaluation standard and guide future method development, including recently emerging foundation models, to advance our understanding of regulatory interactions in longitudinal, multimodal single-cell data.

## Main

Single-cell multiomics technologies have emerged as powerful tools to unravel the complexity of cellular processes. They allow simultaneous measurement of multiple molecular modalities such as DNA accessibility, RNA expression, and protein levels in individual cells. Technologies, such as 10x Multiome^1^ and CITE-seq^2^, are being used extensively to generate novel datasets^3–5^. These paired measurements in a large number of cells enable in-depth characterization of cell states or inference of gene regulatory mechanisms^6^. Regulatory network inference algorithms predict the expression of one gene from all other genes^7,8^ and simple models such as GRNBoost2 have recently been shown to outperform more complex methods^9^. However, current regulatory network inference methods show limited generalizability across conditions and network completeness^9,10^, underscoring the need for novel approaches to improve applicability. A promising, but underexplored direction is to reframe the problem as a cross-modality prediction task by predicting gene expression from chromatin accessibility or surface protein expression from gene expression data. For accurate, generalizable cross-modality prediction a model should learn the biological regulatory processes and identify important features for cross-modality prediction in specific cell states.

Several methods have been developed to achieve cross-modality prediction. MultiVI^11^, scButterfly^12^ and BABEL^13^ are specifically designed to predict gene expression from chromatin accessibility, while totalVI^14^, scPENN^15^ and cTP-net^16^ predict surface protein levels from gene expression. Other frameworks such as Seurat^17^, scVAEIT^18^, scArches^19^ and GLUE^20^ embed any multi-modal data and can infer missing modalities. Furthermore, we, the Open Problems in Single-cell Analysis project (www.openproblems.bio)^21^, hosted our first multimodal single-cell data competition in 2021, where participants were asked to predict modalities in a bone marrow dataset. Remarkably, the top performing solutions of the teams Guanlab^22^ and LS-Lab^23^ were recently shown to perform on par or outperform other methods in an independent benchmark^24^, highlighting the ability of competitions to provide new state-of-the art solutions. However, existing modality prediction benchmarks do not cover the temporal variation in biological regulation. Regulatory interactions between modalities are rarely static, frequently changing during cellular development or upon perturbation. Consequently, models trained exclusively at single time points may capture local cell state transitions, but fail to capture global distribution shifts during biological processes such as cell differentiation or disease progression and cannot reliably extrapolate future biological states^25,26^.

To address these challenges, we generated the largest longitudinal single-cell dataset capturing chromatin accessibility, RNA, and surface protein expression and organized a Kaggle competition that attracted 1,600 participants with various backgrounds. They were challenged to solve two tasks of predicting RNA levels from open-chromatin regions (Multiome task) and surface proteins from RNA levels (CITE-seq task). Our competition set-up followed the Common Task Framework^27^, which has proven to be successful in a wide range of machine learning tasks. It consists of a publicly available training data set, a private test data set only released after the end of the competition and a predefined evaluation metric used for the ranking of solutions. The longitudinal nature of the dataset provided us a natural test set drawn from a distinct distribution of gene-gene relationships. Methods were provided early time points for training, and evaluated at a later, unseen time point using Pearson’s R as an evaluation metric.

Here, we describe the outcome of the competition and the results of a comprehensive follow-up study that contextualizes results from a methodological and biological perspective. We identify successful modeling approaches and perform an extensive ablation study of the top performing methods to identify the minimal set of decisions required for outstanding performance. We further examine validation strategies, which enable selecting robust and generalizable solutions. Furthermore, we evaluate how leveraging prior biological knowledge facilitates modality prediction, identify research gaps where it doesn’t, and demonstrate how top performing models learn regulatory interactions between modalities via feature importance analyses. We envision that this study will set standards for the evaluation of modality prediction methods, provide guidance to method developers, and ultimately impact our ability to infer regulatory interactions from single-cell multimodal data.

## Results

### Advancing Multimodal Time Series Modeling through an Open Problems Competition

At the 35th Neural Information Processing Conference in November 2022, we organized the “Open Problems - Multimodal Single-Cell Integration” competition. The competition was hosted on Kaggle^28^. Participants were challenged to predict omics profiles across modalities in longitudinal single-cell data. This design was set up so that successful approaches would have to model covariation between modalities across biological contexts (time points). This encouraged methods to indirectly capture gene regulatory relationships. To provide a new training and evaluation dataset for this competition, we generated the largest longitudinal single-cell data set at the time capturing three modalities, which is still the largest of its kind to date (**Supplementary table 3**). It comprises over 280,000 CD34+ cells extracted from blood samples of 4 donors, differentiated *in vitro* for 10 days and sampled at 5 time points (**Fig. 1a**). Each sample was profiled jointly for snRNA-seq and chromatin accessibility (10X Multiome) as well as scRNA-seq and protein abundance (CITE-seq), resulting in two complementary datasets that measure 23,418 genes and 228,942 peaks in 161,868 cells and 22,085 genes and 134 proteins (or antibody-derived tags, ADTs) in 119,191 cells, respectively (**Fig. 1b**). To enable modeling of a longitudinal process, we selected hematopoiesis via in vitro differentiation as a model system, characterized by dynamic changes in cell type composition over time, including a decreasing proportion of hematopoietic stem cells and increasing proportions of progenitor cells for hematopoietic lineages (**Fig. 1c**, **Supplementary Fig. 1**).

**Figure 1:**
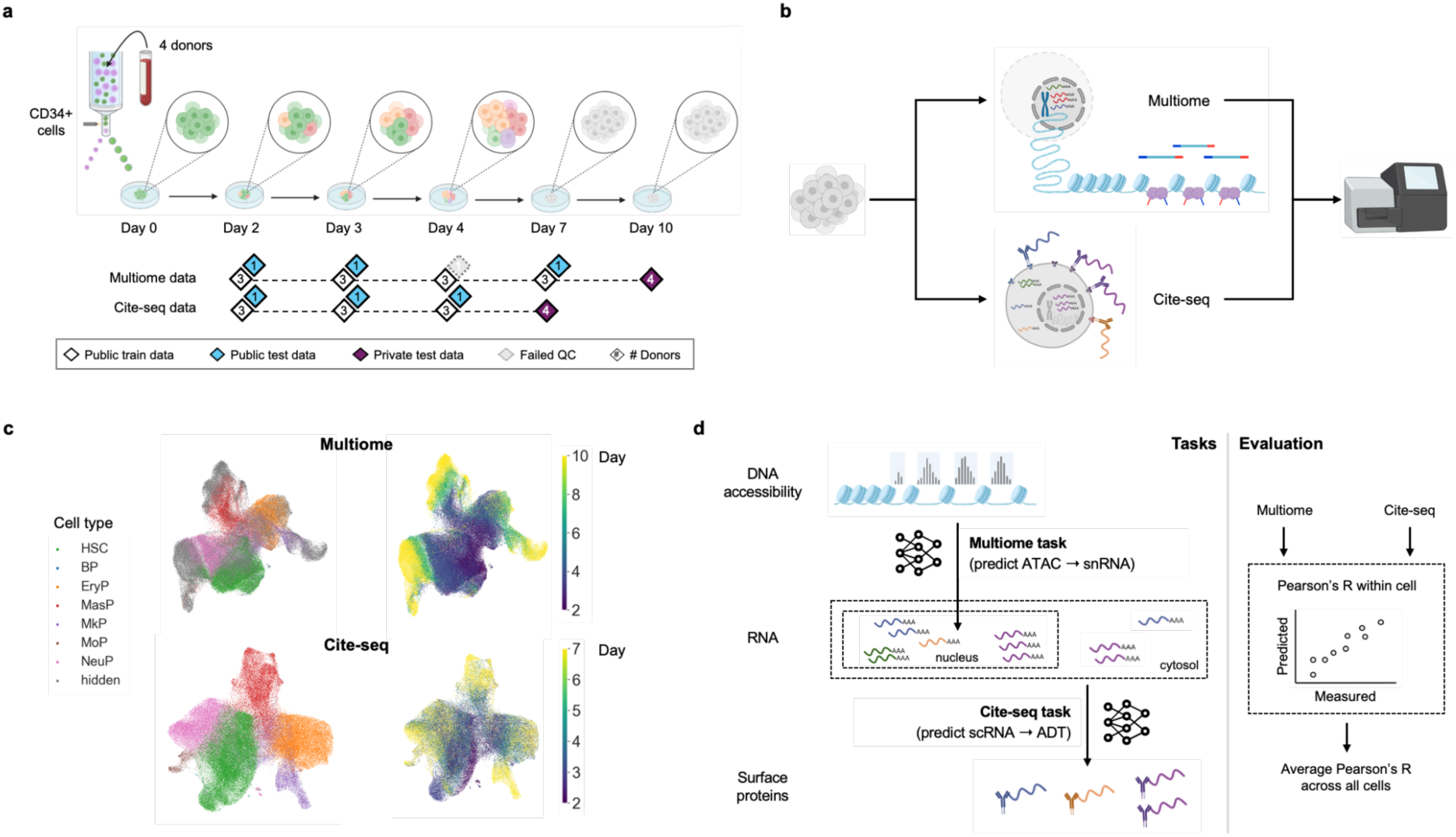
Competition setup, dataset and tasks. **a**, Schematic diagram of the dataset design and train-test splits. CD34+ cells were extracted from blood samples of 4 donors, plated and differentiated for 10 days. For both data modalities, a public train-test split was defined as one of the four donors across the first time points, while the private test set consisted of all four donors at day 7 for the CITE-seq data and day 10 for the Multiome data. Numbers in the diamonds indicate the number of donors in a set. **b,** Schematic diagram of the modalities profiled in each dataset. Two parallel datasets were measured via Multiome profiling measuring scATAC and snRNA, and CITE-seq profiling measuring scRNA and a panel of 134 surface proteins. **c,** UMAPs of the scRNA-seq and snRNA-seq data from the Multiome and CITE-seq data, respectively, showing the differentiation from hematopoietic stem cells (HSC) to mast cell progenitors (MasP), megakaryocyte progenitors (MkP), neutrophil progenitors (NeuP), monocyte progenitors (MoP), erythrocyte progenitors (EryP) and B-cell progenitors (BP) over time. **d,** Schematic diagram of prediction tasks in the competition that represent the principal flow of information from DNA to RNA and proteins. Predictions were evaluated by Pearson’s R between the predicted and true target modality profile across all cells. ADT: antibody-derived tag.

To model the flow of genetic information from DNA to RNA to protein, we asked participants to predict gene expression levels from DNA accessibility profiles (Multiome task) as well as surface protein levels from gene expression (CITE-seq task). Both tasks followed the Common Task Framework^27^. We evaluated submissions based on average Pearson’s R between predicted and true target modality within each cell (**Fig. 1d**). This way, we choose a metric invariant to the scale of predictions, learning from our previous competition at NeurIPS 2021^29^. During the competition phase, submissions were assessed using a publicly available test split, which included one of the four donors across the training time points. The final score was determined using a private test split, which consisted of all four donors on a held-out later time point from the same experiment (**Fig. 1a**). While temporal modeling is not explicitly evaluated in this setup, models sensitive to time-resolved changes are expected to perform better due to shifting cell type proportions and cellular states over time. To prevent information leakage between tasks, we evaluated predictions in the Multiome task at a later time point (day 10) than in the CITE-seq task (day 7). This ensured that the input modality in the private CITE-seq data did not closely resemble the target modality in the Multiome task. Additionally, we removed cell type labels from the private test split of the Multiome task (**Fig. 1c**) (see Methods). In summary, this competition was designed to foster the development of innovative methods for cross-modality prediction in longitudinal single-cell data while promoting models that learn underlying regulatory relationships across modalities.

### The competition accumulated know-how across disciplines and advanced state-of-the-art

The competition, which ran on Kaggle from August to November 2022, saw over 27,000 submissions from 1,602 competitors (**Fig. 2a**), making it the largest competition in single-cell data analysis at the time and to the date of writing this paper. A post-competition survey filled out by 30 participants revealed varying experience in single-cell data analysis and varying domain expertise (life science and other fields; **Fig. 2a**, **Supplementary Fig. 2**). During the competition, participants engaged in active discussion, with over 100 threads on the competition discussion board^30^. Topics ranged from data preprocessing to model architecture and domain-specific interpretation. Several top-performing teams consisted of members combining expertise from machine learning and the life sciences. Furthermore, post-competition community efforts, such as webinars and collaborative forums, indicated sustained interest in interdisciplinary exchange and further research development beyond the scope of the competition^31,32^. Thus, our competition supported the exchange of domain knowledge and a wider machine learning know-how.

**Figure 2:**
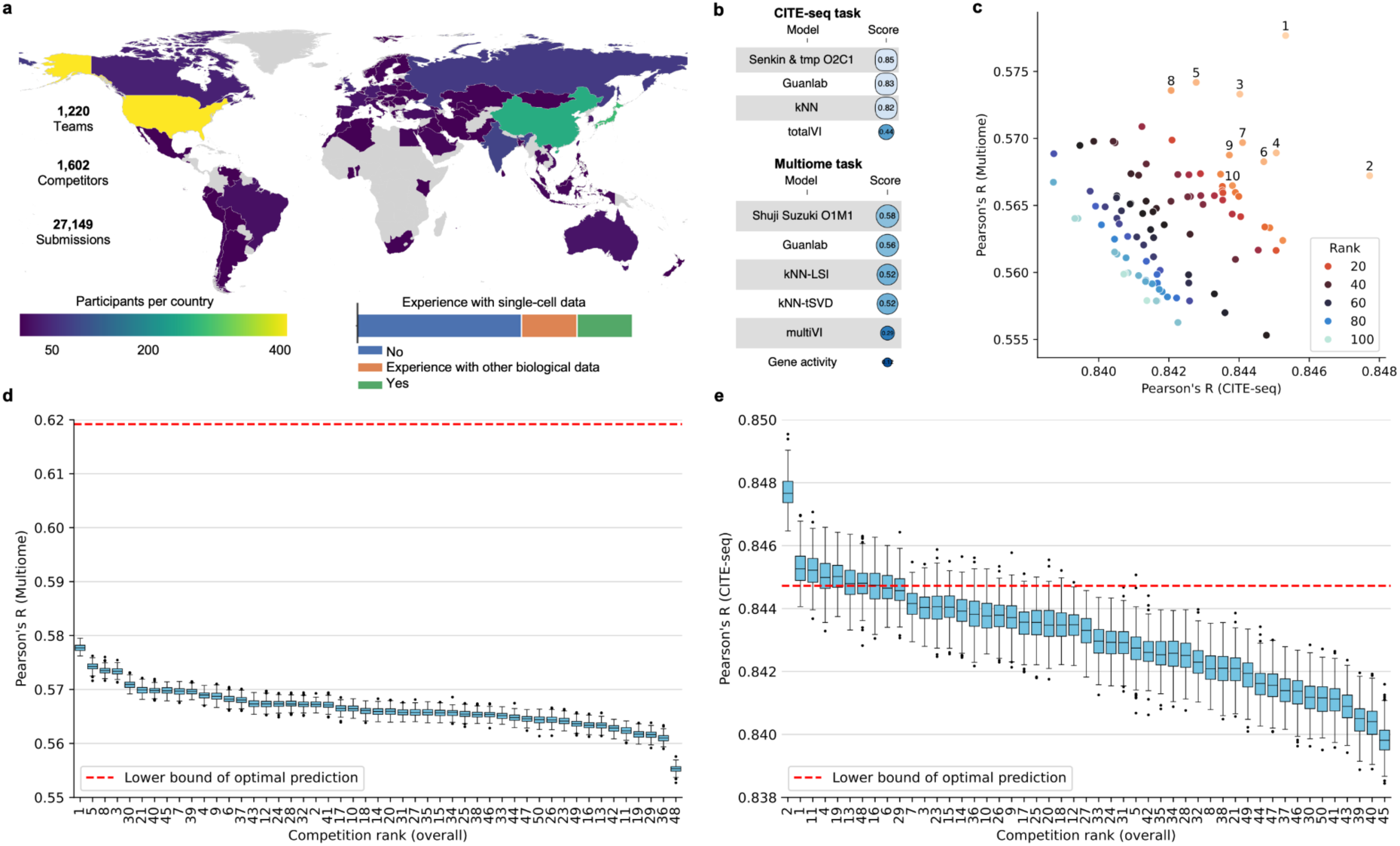
Global participation and highly performant models for the Multiome and CITE-seq tasks. **a**, Map displaying the country of origin of competitors and a survey question results. It shows remarkable participation from all continents and across the skill sets. **b,** Benchmark of Baseline and state-of-the-art prediction models evaluated by Pearson’s R. Top solutions for each task outperform commonly used modality prediction models. **c,** Performance of top 100 solutions at the Multiome and CITE-seq task. Ranks of the leading 10 solutions according to the total score are additionally labeled with the corresponding numbers. This panel shows how the overall winning solution performed specifically well in the Multiome task while the second rank excelled in the CITE-Seq task. **d-e,** Robustness assessment of the ranking and overall performance. Boxplot displaying Pearson’s R values for bootstrapped predictions of the top 50 ranks for the Multiome and CITE-Seq task. The red dashed line represents the lower bound optimal prediction derived from a KNN regression for each cell on the private test data. For both tasks, the winning solution robustly shows superior performance. Overall, the CITE-Seq task was solved specifically well compared to the Multiome task with the top solution outperforming the lower bound optimal prediction.

The final competition ranking was computed by averaging the Multiome and CITE-seq task performance. Competitors used a wide variability of approaches with different preprocessing of the data, modelling and validation decisions, often unconventional for scientific literature. One benefit of the Kaggle competition platform is that winners are required to provide code to reproduce their submissions. After the competition ended, we used these assets to re-evaluate submissions per modality (see **Supplementary table 4** for ranks and **Supplementary table 5** for scores per modality) to investigate which submissions performed best for each task. We found that different competitors excelled in different tasks suggesting that prediction models are best developed for a specific modality (**Fig. 2c**). The overall competition winner *Shuji Suzuki* created the best model for the Multiome task (henceforth called model O1M1; overall rank 1, Multiome task rank 1), and the second place team by *senkin13 and tmp* outperformed other competitors in the CITE-seq task (model O2C1). As the competition was run with both a public and a private test set, we could validate the performance robustness of suggested solutions. Although some solutions had a different performance on the test sets, indicating possible overfitting to the public test set, leading solutions were overall robust. Among 12 solutions that won a golden medal (the top 0.1% of all submissions), 7 kept similar performance for both public and private test sets. For example, the second rank solution moved from first to second place between public and private test set evaluations. Given that the private test set included all samples at an unseen time point and that the ground truth data for 1 donor from the public test was unknown to participants, this suggests good generalizability of the underlying model across time points and donors.

To assess how significant the performance differences between methods are, we evaluated the robustness of the competition results to changes in test datasets, leveraging the availability of all the submissions on the Kaggle platform. Bootstrapping predicted values of the target modality showed that the top rank model for each task robustly performed better than the following ranks (**Fig. 2d-e**). However, when comparing the performance of worse performing models, the rank robustness becomes less pronounced, suggesting that only top performers were significantly better than other methods. We assessed how close the proposed models come to optimal performance by estimating a lower bound optimal prediction computed with full data leakage (see Methods). In the Multiome task, the top solutions did not reach the lower bound optimal prediction, suggesting substantial room for improvement in this task (**Fig. 2d**). In contrast, top performing solutions in the CITE-seq task outperformed the estimated lower bound optimal prediction (**Fig. 2e**). The predictions were uniformly good for all surface proteins in our data with the worst Pearson’s correlation score between true and predicted values being 0.803 (**Supplementary Fig. 8**). These results suggest that top performing models can provide highly accurate imputation of surface proteins and capture complex regulatory relationships between RNA transcription and protein expression.

We next investigated whether top solutions in this competition could outperform approaches commonly used in multimodal single-cell data analysis and state-of-the-art modality prediction models (**Fig. 2b**). We reimplemented baseline models (gene activity scoring^33–35^, KNN regression) and current state-of-the-art approaches represented by the top models from our previous NeurIPS 2021 competition^6^. Baseline methods performed worse than more complex methods, especially for gene expression prediction from chromatin accessibility. Gene activity scoring and kNN regression reached Pearson’s R of 0.12 and 0.52 correspondingly for the Multiome task. The previous state-of-the-art method achieved a Pearson’s R of 0.56, but was surpassed by the top 2022 competition model (O1M1) which reached a Pearson’s R of 0.58. In the CITE-seq task, a baseline kNN regression reached R of 0.82. The top model from our 2021 competition benchmarked at R = 0.83, but was surpassed by the best 2022 competition model (O2C1) which achieved R= 0.85 (**Fig. 2b**). Overall, the above evidence suggests that top performing models in our competition produce highly accurate predictions, especially in the CITE-seq task, and outperform the current state-of-the-art.

### Successful solutions use neural network models and extensive preprocessing of the data

To investigate what distinguishes top performing solutions from less successful ones, we collected publicly available information about modeling approaches, validation strategies, and metadata usage from 15 source code submissions and 23 discussion threads shared on the Kaggle platform (**Supplementary table 5**). Survey results highlighted that neural network-based approaches were particularly popular among top-performing competitors and that a large range of model complexities from 3-11 layers with 128 to 1024 dimensions per layer was explored. Furthermore, half of the responses indicated that cell type labels were used as part of preprocessing or predictions in the CITE-Seq task (**Supplementary Fig. 2**). The winner of the competition exclusively used neural networks (NN), while the rank 2 and 3 models combined Light Gradient-Boosting Machine (LGBM)^36^ and Catboost^37^ models with NNs. Almost all competitors combined outputs of different models in an “ensembling” strategy^38^. To aggregate model results, some competitors used simple averaging (e.g., rank 1 and 5), while others weighted different predictions (e.g., ranks 2-4) either using a linear model to learn weights that maximize the validation score or using weighting guided by intuition. The rank 2 submission leveraged a more sophisticated approach to ensembling that included training a meta-model based on the predictions from other models.

The winning solution (labelled O1M1) by Kaggle user “*Shuji Suzuki*” used a neural network architecture with extensive preprocessing of the input data. It incorporates a multi-head prediction structure and averages predictions from different heads of the model (**Fig. 3a**, see Supplementary information for a detailed description of all models and preprocessing strategies). This approach ranked 1st in solving the Multiome task (**Fig. 2d**) and achieved 2nd place in CITE-seq prediction (**Fig. 2e**). Unique features of this solution included the prediction of residuals for inverse truncated singular value decomposition (TSVD) in the Multiome task, and using biological prior knowledge from the Reactome database^39^ to select genes related to target proteins for the CITE-seq task.

**Figure 3:**
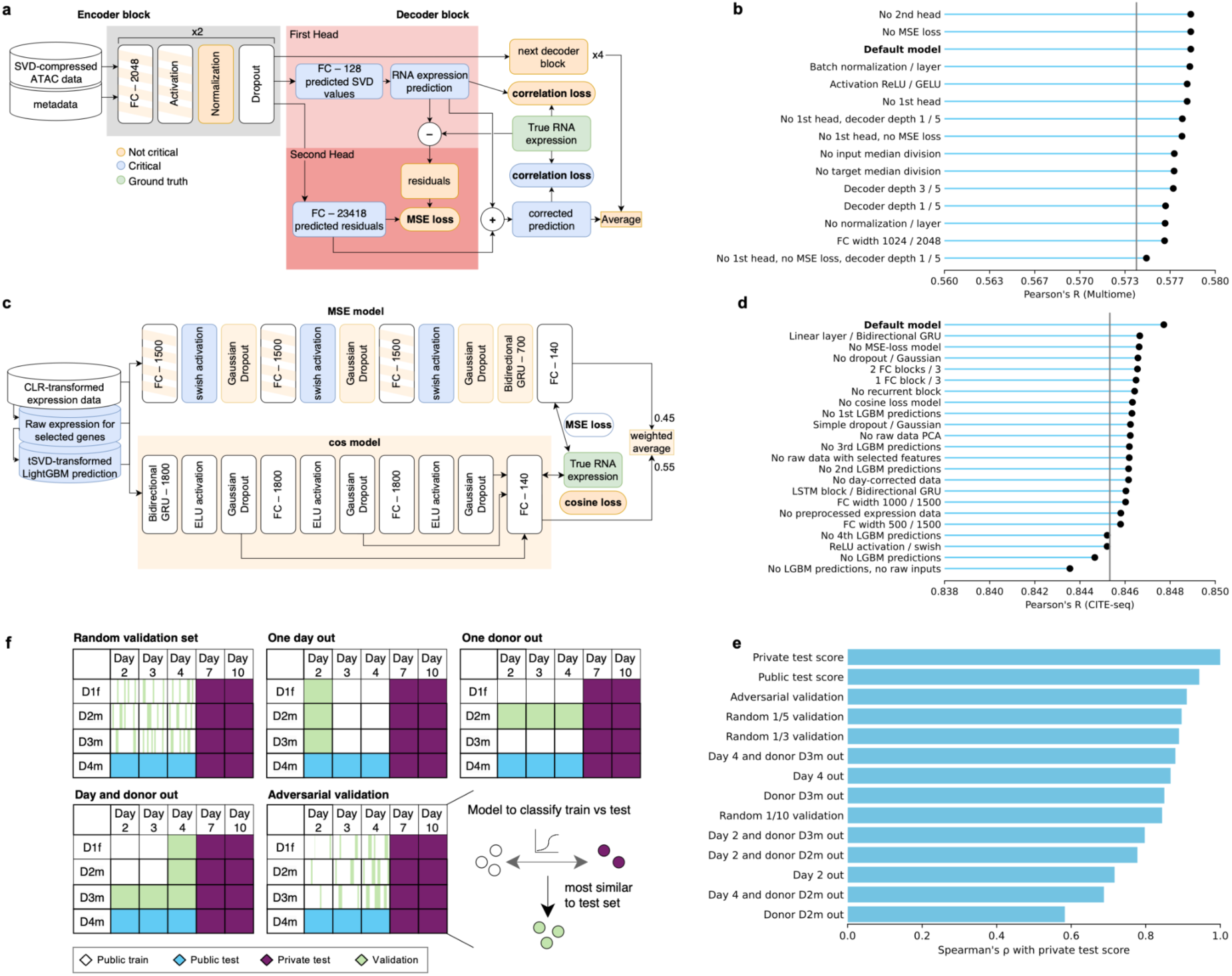
Model architectures of top performers, ablation study and validation strategies analysis. **a**, Architecture of the best Multiome prediction model from place 1 solution (O1M1). Orange blocks are not important for model performance according to our ablation studies and can be removed without a significant reduction in Pearson’s R. Striped orange blocks can be simplified to keep a similar score. Light-blue blocks are crucial for model performance. x2 and x4 for encoder and decoder parts mean that identical blocks are repeated 2 and 4 times, respectively. **b,** Ablation study results for the model from **a)** showing score on the private test set for model variants. “/” in the model names should be read as “instead of”, followed by the value in the original model. The scale matches the spread of scores for top 49 models on Fig. 2d. The grey line marks the score of the second best model for this task. **c,** Architecture of the best performing CITE-seq model from place 2 solution (O2C1). The coloring scheme is identical to a). **d,** Ablation study results for the model from c), presented as in b). The scale matches Fig. 2e**. e,** Cross-validation strategies visualization. Light green illustrates cells taken as a validation set to test the performance of different models. In the random validation strategy, a fixed proportion of cells is taken with an equal probability of every cell to appear in the validation set. In day- or donor-out strategies, all cells from a particular day, donor, or their combination are left out for testing. Adversarial validation requires training a classifier to predict if a cell comes from a training or testing set. The fixed number of cells from public training data with the highest predicted probability of coming from the test set are later used for the model validation. **f,** Validation strategies quality calculated as an agreement of model ranking by a score on a validation set and by score on the private test set.

The second-place team (O2C1) *senkin13 and tmp* combined inputs from multiple preprocessing methods that were fed into LGBM models and into 2 neural networks with different loss functions and slightly different architectures further aggregating results by a weighted average (**Fig. 3c**). This solution holds the first place on the public test evaluation, showing consistently good performance on both test sets, and particularly excelled in the CITE-seq task, outperforming all other approaches by a wide margin (**Fig. 2e**). Notable features of the second place approach are the use of multiple preprocessing approaches for the input data, including CLR transformation of RNA expression data, a custom normalization approach (combining mean expression scaling factors, a square root transformation, and z-scoring per gene), selecting the most correlated genes for each target, and TSVD transformation (**Supplementary Table 2**). Datasets preprocessed in various ways were used to obtain predictions from gradient boosting models, and their outputs were used as an additional input to neural networks (see Supplementary for an extensive description).

The third-place solution (henceforth: O3), by Kaggle user *Makotu*, relied heavily on ensembling numerous models and datasets. 20 models were trained for the CITE-seq task, including 18 neural networks and 2 Catboost models, and 26 neural networks were used to obtain prediction in the Multiome task. The input datasets included SVD-transformed normalized and raw counts data, selected features with the high correlation to the targets, word2vec-transformed data^40^, cell type proportions and aggregated data per cluster or donor. Besides various preprocessing, a unique feature of this solution was an adversarial validation strategy used to locally evaluate the quality of the models (see the section below).

### Winning models can be simplified while preserving prediction quality

To gain insights into best practices for predicting RNA from chromatin accessibility and surface protein abundance from RNA expression, we deeply investigated the 3 highest ranking solutions, using the code and solution descriptions published on the Kaggle platform. We performed ablation studies of the winning models to identify critical features of their modeling approaches. Specifically, we investigated the effects of preprocessing, loss functions, model architecture, model width and depth, activation functions, normalization techniques, and data preprocessing (**Methods**). We selected O1M1 and O2C1 models for their impressive performance on the Multiome and CITE-seq respectively, and O3 model for a good overall performance and an outstanding number of applied input data preprocessing methods.

For the Multiome task, we ablated the M1 model to identify modeling components that are critical for preserving performance. We discovered that ablating most of the individual parts, including MSE loss for the residuals prediction from the second head, normalization layers, decoders’ first heads predicting SVD-compressed values of expression or second heads predicting the residuals of transcriptome restored from predicted SVD, and even leaving only 1 instead of 5 decoder blocks kept the performance very similar to the original model (**Fig. 3b**). Only the combination of ablations, namely leaving 1 of 5 decoder blocks, removing MSE loss for residual prediction, and the entire first head, resulted in dropping performance down to the level of the 8th, 5th, and 3rd place submissions for the Multiome task (**Fig. 2d**). These results show that the top-performing prediction model for the Multiome task can be significantly simplified while keeping outstanding performance.

To identify critical model components for the CITE-seq task, we ablated the O2C1 and O3 models, thus focusing on the best solution for this task, and on a less performant approach with rank 13 for the CITE-seq prediction. For the O2C1 model, which averaged predictions of 2 neural networks (**Fig. 3c**), the aggregated prediction resulted in a better performance (Pearson correlation with the private test set Pc=0.8477) compared to single model predictions (Pc=0.8463 and Pc=0.8466 for MSE loss and cosine loss models respectively), while individual models had similar scores. We focused our ablation study only on the model trained with MSE loss because of its comparable simplicity. The least important parts were bidirectional GRU layer, dropout blocks and 3 consecutive fully connected layers as removing or simplifying them resulted in comparable or even slightly better prediction compared to the original MSE loss model. Elements crucial to model performance included the diverse preprocessing of input data and the predictions of the LGBM models used as additional input features to NNs. When LGBM predictions and raw counts-based data are not used, the model performance dropped to P_c_=0.8436, making it worse than the top 100 solutions. These results indicate that using NNs to correct predictions of simpler models is a powerful approach, which is in line with the prediction strategy of the O1M1 model, which relies on correction of the data obtained with inverse SVD transformation.

The O3 CITE-seq prediction was based on a complex ensemble of 20 models trained on different combinations of 27 variants of the data with diverse preprocessing (See Supplementary information) and was a perfect test case to evaluate the effect of ensembling. We tested the predictions of the ensemble with one or two models ablated, as well as the predictions of individual models on public and private test sets (**Supplementary Figure 4**). When a single model was used for prediction, private test scores were significantly lower than in ensembles of 18, 19 or all 20 models, often pushing the submission out of the top 10. Interestingly, one of the best performing individual models used the simplest preprocessing approach containing 84 selected genes as well as 128 SVD components of log-normalized expression data excluding these 84 genes. The NN model trained on this simple dataset alone would have ranked 8 in the competition, indicating that complex preprocessing is not necessary for good performance.

We additionally created ensembles of 4 models with the best private test scores and 4 models with the worst private test scores. While this would be an unrealistic scenario at the competition because the ground truth score was unknown, this experiment demonstrates an interesting effect. The average prediction of 4 worst individual models predicted the data better than most single-model predictions, and the ensemble of 4 best models received an even better score than when all 20 models were used. Thus, ensembling is a powerful strategy even when the individual models are not the best performers, but adding too many models in the ensemble can lead to a drop in prediction quality.

### Validation strategies have a major impact on prediction robustness

The competition structure with public and private test sets forced competitors to create models that generalize to new samples and time points in an out-of-distribution scenario. Competitors evaluated and optimized for this generalization capability using different cross-validation approaches. Competitors labeled part of the training data as a validation set, trained different models on the rest of the data, tested performance on the validation sample, and submitted only the best-performing model (or more frequently, an ensemble of the best models) retrained on the whole dataset. This approach relies on the assumption that the validation score is monotonically related to the private test score. To determine the best practices for model development in our competition, we investigated the validation strategies used by competitors by systematically evaluating the effect of exchanging validation strategies for top performing models.

The participants of the competition utilized several validation strategies. Following the known structure of the test sets, competitors often used the leave-day-out or leave-donor-out strategy, evaluating model quality on the data from a day or donor unseen during training or hiding both one day and one donor from a training set. This approach was particularly popular among the top positions of the leaderboard. Solutions at places 1, 5, 6, 7, and 8 all used validation split by day and donor; the solution that took the 2nd place leveraged validation on an unseen day, and the solution at place 4 was obtained using validation on an unseen donor. A female donor was often not included in a validation set because participants uncovered^41^ that the donor in the private test set was male and assumed that validation on another sex would not be reliable^42^. While randomly splitting the data into 3 or 5 folds was a common validation strategy, no method in the top 15 used this approach for both tasks, suggesting that random validation has a worse ability to find generalizable models. The 3rd and 41st place solutions used a so-called “adversarial validation” technique^43^, where a classifier was trained on all available transcriptomics data to label each cell as belonging to the public train set or a private test set. Cells from the train data misclassified as belonging to the test set were used for validation. This strategy is designed to identify cells in the public train split that are closest to the private test set and enable robust local evaluation of models with known ground truth. We noted other approaches for validation as well but did not investigate them in detail as they were associated with poorer performance. It is worth mentioning that the validation strategy is unknown for 11 of the top 25 solutions (all ranked between 9 and 25).

To assess the effect of common validation strategies, we systematically compared the ranking of variants of top-performing models by performance on different validation sets. We conducted this analysis only on the CITE-seq prediction models as participants tended to use more complex validation strategies only for this task due to the reduced computational resource requirements. We trained 23 variants of the neural network models from the top 3 participants, tweaking the width and depth of some models, activation functions, normalization approaches, and random seeds (see Methods). We tested 12 validation strategies: 3 random validation sets containing one-fifth, one-tenth, or one-tenth of cells in the training data, 2 leave-day-out sets excluding days 2 and 4 (the first and last days in the training data), 2 leave-donor-out sets with male donors (D2m and D3m), 4 combined leave-day-and-donor-out strategies, and an adversarial validation strategy containing 4000 cells from a train set that were misclassified by a LGBM model as test cells (**Fig. 3e**). Validation strategies were evaluated based on the similarity of the overall ranking of models to the ranking on the private test set used to score the competition.

The best predictors of the private test performance were a score on the public test set (Spearman correlation coefficient ρ=0.944, **Fig. 3f**) and a validation score using the adversarial set (ρ=0.910), followed by validations on random one-fifth (ρ=0.896) and one-third (ρ=0.889) subsets of the data. Leaving day 4, or donor D3m, or the combination of both resulted in an intermediate validation strategy (ρ=0.867, ρ=0.850, ρ=0.879 correspondingly) while using the cells of donor D2m or coming from day 2 resulted in poor generalization of performance (ρ=0.583, ρ=0.716, correspondingly). The success of validation strategies may be explained by their transcriptional or compositional similarity to the private test set (**Supplementary figure 3**). However, similarity to the test set was not in general predictive of validation strategy performance. While the public test set showed the highest correlation with the final ranking, optimizing only this score could result in a poor generalization of performance as illustrated by the fall of the O2C1 solution with the 2nd place on public leaderboard to place 133 on the private test set. Together, these results suggest that the adversarial approach is the most generalizable validation strategy.

### Incorporating biological priors does not consistently enhance model performance

Including prior biological knowledge has been shown to be impactful to improve modeling performance in data-scarce settings^44,45^. Thus, we encouraged competitors to include additional datasets and prior knowledge to improve prediction accuracy. However, top-performing solutions generally did not explore this approach. This leaves it unclear whether this lack of biological priors is due to their inability to improve model performance on this dataset or whether it was simply not tested. To determine whether biological priors could be used to improve model performance, we systematically evaluated the effectiveness of integrating prior biological knowledge using biologically driven feature engineering approaches with a simple baseline model using SVD-transformed input data on a 2-layer neural network (**Methods**).

In the CITE-seq task, we evaluated the inclusion of different protein-protein interaction (PPI) network-derived features hypothesising that they reflect gene regulatory interactions. Features were obtained by propagating the expression levels of interacting proteins for each target protein whose abundance was predicted using random walk with restart (**Fig. 4a**). We tested multiple network sources (STRING^46^ and HumanBase^47^ networks) and propagation depths ranging from zero (no propagation) to five hops. Across all network types and propagation depths, the addition of network-propagated features improved model performance only marginally (0.25%) from ρ=0.8407 (SVD baseline) to ρ=0.8428 (SVD & 0-5-hop combined) and performance was largely unaffected by the choice of network source (**Supplementary Fig. 5**). Models trained exclusively on network features performed worse than those including SVD components, indicating that information contained in the networks is already captured by the transcriptome-derived features. We tested different scales of network features by varying the number of hops in the network propagation from zero (0-hop; no propagation) to five propagation hops using a random walk kernel (**Methods**). 0-hop network features that use only expression levels of genes that match the target protein were the most performant feature that could be added (**Fig. 4b**). Including 0-hop features was indeed also a strategy used by multiple top performing competitors including the best performing CITE-Seq model. However, comparison with bootstrapped results from a similarly performing model (overall rank 49) showed that these differences lie within the range of sampling variation of the test set, indicating a small effect size of adding network-based features.

**Figure 4:**
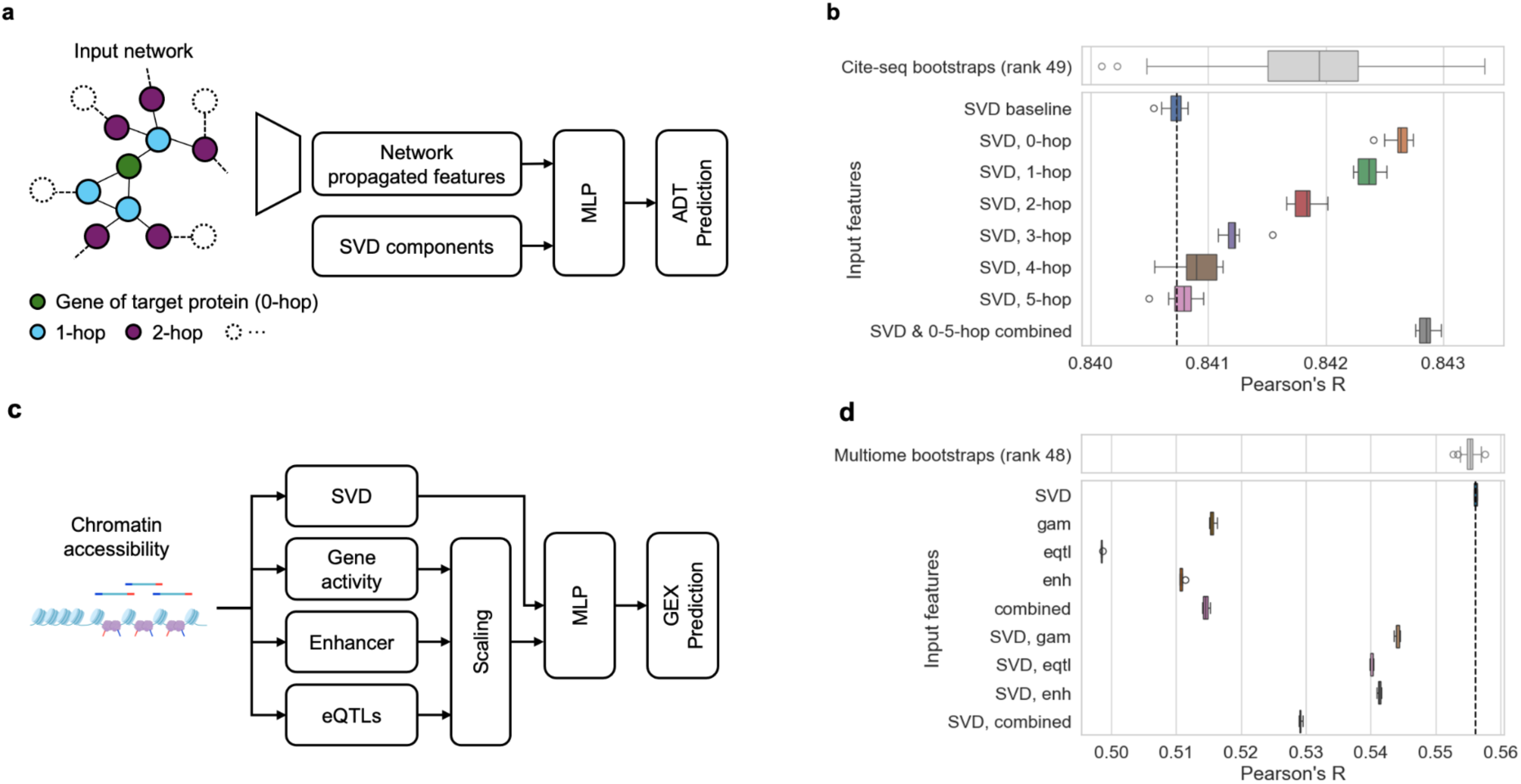
Integration of prior biological knowledge and interpretability of feature attributions. **a**, Schematic drawing of the model to evaluate the integration of protein-protein interaction (PPI) network-derived features in the CITE-seq task. Network-derived features were combined with SVD components to predict surface proteins. **b,** Boxplot showing model performance with varying depth of network-propagated features added to SVD components. It shows that adding 0-hop (direct gene expression of target proteins) and the combined set of features can improve model performance. Above the bootstrapping results of a model with similar performance is shown indicating the scale of performance differences when adding network-derived features. **c,** Schematic drawing of the model to evaluate integration of prior biological information in the Multiome task. Accessibility of the gene body and promoter region (gene activity), known enhancer regions and known eQTLs were added to SVD components. **d,** Boxplot, displaying model performance of predicting gene expression from chromatin accessibility and bootstrapping results of a model with similar performance. In all cases, addition of prior-bio-derived features decreased model performance.

In the Multiome task, we evaluated feature engineering approaches following three known mechanisms of gene regulation and combined these with SVD components (**Fig. 4c**). Specifically, we computed accessibility features in gene bodies and promoters (gene activity matrix, GAM), known enhancer-gene associations from multiple databases^48–51^ and genomic regions of known expression quantitative trait loci (eQTL), whereby higher accessibility is generally associated with increased gene expression. To compare the effect of different biology-informed features independent of the number of features available for each type, we reduced the set of target genes to 1115 genes for which all feature types were available. Surprisingly, we observed that adding biology-informed features decreased performance across feature types on the private test data substantially compared to only using SVD components (**Fig. 4d**). We investigated this decrease of performance by testing how well each feature set can represent cell types - a simplified task to evaluate the information content of the biologically informed features. Principal component regression analysis, which computes the variance associated with different covariates (**Methods**), showed that cell type differences were associated with three times less variance in the three prior knowledge-derived feature sets compared to SVD components (**Supplementary Fig. 6a**). Furthermore, we explored whether the correlation between regulatory features and their corresponding target genes changes during the differentiation process, which could explain decreased model performance at later time points in the private test set. We indeed identified shifts in the correlation structure even within the same cell types, suggesting that direct promoter-gene links become less predictive as differentiation progresses, possibly due to the increasing influence of post-transcriptional regulation not captured by chromatin accessibility alone. This could further explain the insufficient performance of predicting gene expression at a later time point using prior knowledge derived features (**Supplementary Fig. 6b**).

Box 1: Best practices for cross modality prediction

As a result of the deep investigation of the top 3 solutions, a survey of the competitors, and the ablation study, we formulate the following best practices for building a model for the prediction of RNA from chromatin accessibility data or surface proteins from RNA expression data:

- Neural networks show a superior performance compared to tree-based and other classical machine learning models
- Fully connected networks with 1-4 hidden layers perform the best on the current scale of the data
- While complex activation functions, such as swish, may work better in certain models, ReLU activation often leads to a similar performance. We therefore recommend starting from the simplest approaches and then testing more complex modelling decisions.
- Combining different preprocessing regimes of inputs helps to achieve better performance
- Ensembling significantly improves prediction quality and robustness of the results
- Feature selection, either data-driven or based on prior knowledge, significantly helps to achieve good predictions
- Initial prediction with simple models and training a NN model to correct them is a powerful approach
- Models can learn regulatory interactions from the data directly, while effective use of PPI networks or known gene regulatory elements requires more sophisticated modeling strategies than were tested in the competition

### Top performing methods learn biologically relevant regulatory interactions

Deriving biological relevance from top-performing models could offer significant insights into DNA-to-protein regulatory mechanisms. Specifically, within the context of the CITE-seq task (which showed overall better model performance), we assume that a model demonstrating high performance is likely to have captured generalizable RNA-to-protein regulatory relationships. Key predictive features fall into two categories: genes that covary with the target protein due to shared upstream regulation in the given biological context, or genes that regulate the target protein. In the diverse cellular profiles measured across cell types, timepoints, and donors, shared pathways (that indicate shared upstream regulation) are likely activated differently across conditions, making RNA-to-protein regulatory relationships more predictive. This regulation captured by important features for highly performant models is hypothesized to account for the imperfect correlation between RNA and protein levels^52^. Understanding and interpreting these models can therefore offer insights into the underlying mechanisms of protein regulation.

To investigate this hypothesis, we conducted feature attribution analysis using SHAP^53^. The third-place model from Kaggle user Makotu was chosen for this analysis because it required relatively fewer preprocessing steps to revert to the original feature space (genes), ensuring direct interpretability when using feature attribution methods. Given that some models in Makotu’s ensemble clearly outperformed others (**Supplementary Figure 3**), we focused on computing SHAP values for these dominant models.

Overall, we find that genes with high feature importance in the rank 3 model relate to post-transcriptional regulatory signals, confirming our hypothesis. As an example, we run gene set enrichment on the top 20 genes (**Methods**) which are most predictive of CD86 and find post-transcriptional regulation pathways and regulators (**Fig. 5, Supplementary Figure 7**). C1QBP and EIF5A are the 7th and 9th ranked genes, respectively. Eukaryotic initiation factor 5a (EIF5A) promotes initiation of translation and elongation^54^. C1QBP, also known as p32, inhibits splicing factor 2 (SF2) within the nucleus. While C1QBP has been studied in mitochondria, little is known about the nuclear mechanism nor its relation to CD86 regulation, making C1QBP a promising target for further investigation^55^. To ensure that these genes are not just assessed as important for the model because their RNAs are highly correlated with the target protein, we also ranked the RNAs according to their Pearson correlation with the target protein. In contrast with SHAP value analysis that identified post-transcriptional regulators, the top correlated RNAs are enriched for an adaptive immunity response gene set (**Fig. 5**). We reason that this is because CD86 is an antigen presented by monocytes which contributes to adaptive immunity, and these RNAs are correlated because they act in shared pathways. Instead, by looking at regulators obtained from feature attribution scores on a high-performing model, we can identify potential direct regulators of CD86 in hematopoiesis. Thus, top performing models in our competition learned meaningful regulatory interactions to enable accurate prediction of unseen cellular modalities in a new time point.

**Figure 5:**
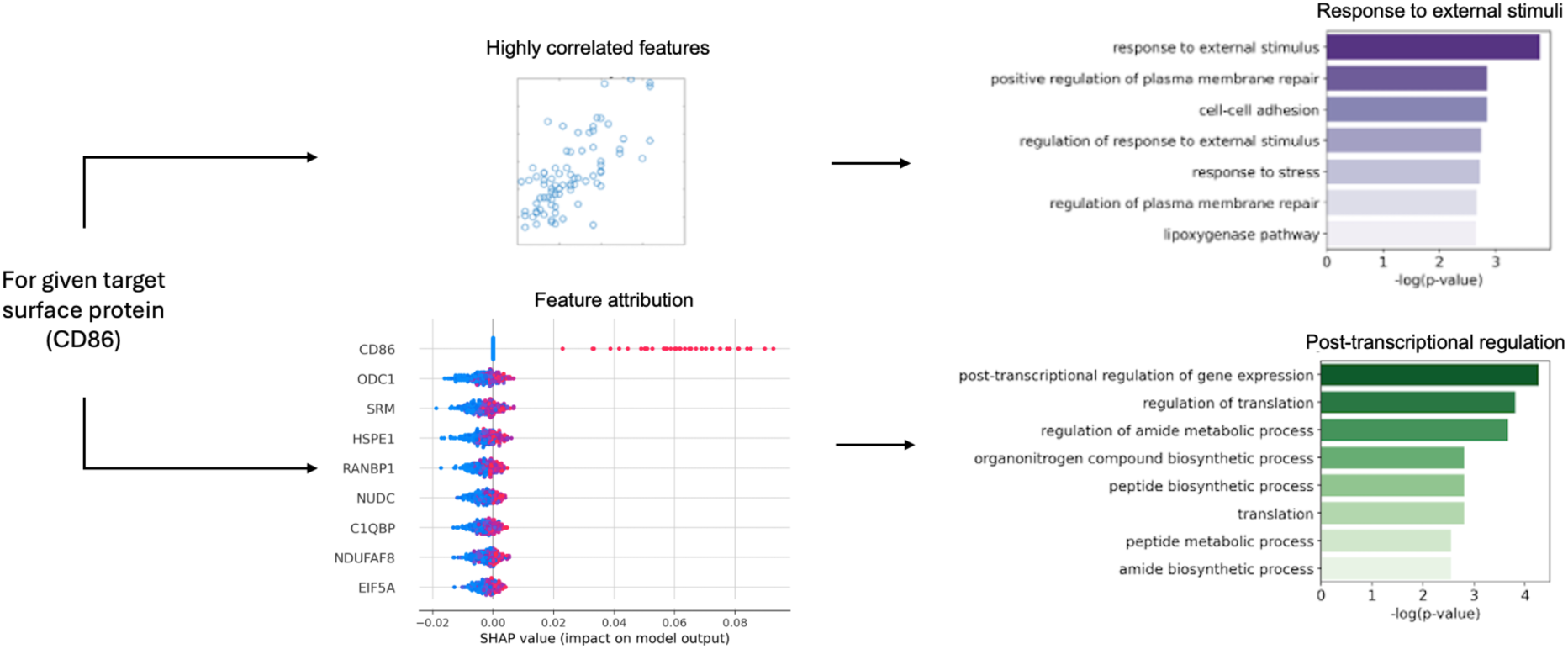
CITE-seq model learns biologically relevant regulatory interactions. Shown are the most important features (highest SHAP values) in the rank 3 solution for predicting CD86 levels compared to highly correlated input features. While highly correlated features are associated with the biological function of CD86, the suggested model learned features associated with post-transcriptional regulation.

## Discussion

We hosted the largest single-cell data science competition to date, asking over 1,600 competitors to predict modality profiles in the direction of the central dogma of molecular biology: predicting RNA expression from ATAC data (Multiome task) and predicting surface protein levels from RNA (CITE-seq task). To power the competition, we generated the largest multimodal longitudinal single-cell datasets from human donors, specifically designed to empower the development of models generalizable to unseen time points and donors, and uncover regulatory mechanisms of hematopoiesis. In follow-up studies, this dataset may also serve as a valuable resource to investigate mechanisms of hematopoiesis with therapeutic relevance. Leveraging over 30,000 submissions, we improved upon existing state-of-the-art approaches for modality prediction, identified best practices for model building **(see Box 1)**, and showed that top performing models learn meaningful regulatory patterns across omics modalities.

Promoting method development via a competition on a single dataset comes with the risk of methods being overly tailored to the set challenge and lacking generalizability. While the solutions proposed during the competition were tuned for the specific dataset and evaluation metric, the competition’s train-test split was specifically designed to measure generalizability: predictions were evaluated across four biological replicates on an unseen time point. Indeed, our benchmark showed that the winning models for the Multiome and CITE-seq task had superior performance compared to the rank 1 solutions from our previous modality prediction competition at NeurIPS 2021. Top performing methods from the NeurIPS 2021 competition in turn have recently been shown to perform on par with best performing state-of-the-art approaches in a recent independent benchmark based on a total of 36 datasets and 6 evaluation metrics^24^, supporting the generalizability of our competition setup. Furthermore, we hypothesize that models that learn interpretable, biologically relevant features should be more generalizable to new contexts, as the underlying regulatory mechanisms they capture are likely to be invariant across conditions, including different time points. When assessing feature attribution scores in the CITE-seq task, we observed an enrichment of features associated with post-transcriptional regulation, which indicates that models learned biologically meaningful, and therefore generalizable, patterns in the data.

Models in the CITE-seq task achieved particularly strong performance, reaching a Pearson correlation of 0.848 for the unseen time point and exceeding our estimated lower bound of optimal performance. Accurate predictions of surface proteins imply that in future CITE-seq experiments in well-characterized tissue types such as HPSCs might be replaced by prediction of protein levels from RNA expression data, unless cells are strongly perturbed relative to conditions captured in available training data. In the Multiome task, the highest Pearson correlation of 0.578 did not reach our lower bound estimate of optimal performance. This performance difference may be connected to the higher dimensionality of the Multiome data limiting both the number of hypotheses that competitors could test in a given time due to computational limitations and the set of tools that could scale to this number of features. Consequently, predicting RNA expression from chromatin state remains a challenging problem. Future improvements may require incorporating additional epigenetic features beyond chromatin accessibility, such as methylation and histone modifications. Regarding protein prediction, we anticipate that standard CITE-seq panels might become less central as novel technologies emerge that can capture the full proteome. Such advances will also enable the design of future competitions focused on predicting the entire proteome.

Our competition enriched the single-cell field with wider machine learning expertise. Participants often used preprocessing techniques unconventional for single-cell data analysis^56^, such as dividing expression values by non-zero median or TSVD-based imputation. Another innovation was adversarial validation technique enabling selection of generalizable models and their hyperparameters based on the cells most similar to test data. Beyond competition setup, adversarial validation can be used when a target sample for prediction is known. With respect to model architectures, fully connected neural networks represented the vast majority of implemented model architectures suggesting that this is currently the best model class for modality prediction. Ensembling predictors is a common approach to improve generalizability in machine learning, however, with few exceptions^57,58^, it is not commonly used in single cell data science. Our competition results suggest that ensembles of models with varying input features or model architectures, can increase performance and robustness of the predictions while remaining interpretable as we showed in SHAP values analysis. Remarkably, in our ablation studies, we showed that top-performing models in both tasks could be simplified substantially, while maintaining rank 1 performance. We therefore provide lightweight yet powerful models to users, and understanding of crucial model components for method developers.

Despite the more than 30,000 solutions submitted during the competition, two modeling directions remained underexplored in the competition and warrant further investigation: the incorporation of longitudinal information and the use of prior biological knowledge. Most competitors, including the top performers, considered the longitudinal aspect of the data only in the validation scheme (and sometimes during data preprocessing), but not in the model itself. This implicitly assumes a static relationship between modalities across time points, ignoring known temporal dynamics. Perhaps a more successful approach would leverage trends of chromatin accessibility, mRNA and protein levels during differentiation trajectories, including time lags between modalities that have been observed and used to compute chromatin potential^51^ and multimodal velocities^52^. Additionally, cell type information was rarely used as direct model input, assuming that gene expression and protein translation mechanisms are not dependent on a cell type or that this information can be implicitly learned from the data. The former assumption is not true in a general scenario^61,62^, but might hold for our dataset coming from one tissue. Interestingly, prior biological information was rarely considered by competitors. Several databases exist that link biological features across modalities, for instance eQTLs or pQTLs to link genomic regions to gene expression levels or protein-protein interaction networks^46–48,50,63^, which could give prior information on co-regulated proteins to inform predictions. We found that the inclusion of this information is not always straightforward and can even worsen model performance in our own efforts. Thus, either these resources are not generalizable to our specific tissue, effect sizes are too small, feature interactions are already sufficiently captured by the data, or modeling approaches for including biological prior information must be improved. Here, we see potential for advancing methods that include gene regulatory networks for the purpose of modality prediction.

As competitions will continue to be a powerful strategy to facilitate method development, we reflected our set-up and gained knowledge for future competitions. While we learned from our previous competition that scale-invariant metrics such as Pearson’s R are important to fairly evaluate predictions on unseen batches^23^, our choice of metric still has room for improvement. Specifically, the sparsity and high dimensionality of the data led to quite small absolute differences in the evaluation metric, which could be improved by giving more weight to less sparse features in the target modality.

Engaging a record number of 1,600 participants, our Open Problems in Single-cell Analysis competition has demonstrated the impact of encoding biologically relevant open single-cell challenges in a quantitative manner to engage researchers across domains. This model of promoting innovation has seen increasing focus in recent years, and we believe our competition can serve as a useful blueprint for future competitions across computational biology. We have advanced modality prediction methods while implicitly learning gene regulatory patterns in the process, exploring a new avenue for multimodal inference of gene regulation that relies on improving modality prediction models while using comparatively simple feature importance approaches. Our best practices for modality prediction, together with our unique longitudinal, multimodal dataset, will provide a foundation for future innovations, extending beyond modality prediction and inferring gene regulation to also enable method development in temporal modeling. Taken together, we envision that the insights from this competition will set standards and drive progress in modeling gene regulatory mechanisms via model interpretation, facilitating a deeper understanding of complex biological systems.

## Methods

### Data Acquisition

The dataset comprises single-cell multiomics data collected from mobilized peripheral CD34+ hematopoietic stem and progenitor cells (HSPCs) isolated from four healthy human donors. The cells were bought from AllCells and more information about the cells can be found under https://allcells.com/research-grade-tissue-products/mobilized-leukopak/.

Measurements were taken at five time points over a ten-day period. During this time, cells were cultured with StemSpan SFEM media supplemented with CC100 and thrombopoietin (TPO) and incubated at 37°C. Media was changed every 2-3 days. No additional media supplements were added to the cell culture conditions.

From each culture plate at each sampling time point, cells were collected for measurement with two single-cell assays. The first is the 10x Chromium Single Cell Multiome ATAC + Gene Expression technology (Multiome) and the second is the 10x Genomics Single Cell Gene Expression with Feature Barcoding technology technology using the TotalSeq™-B Human Universal Cocktail, V1.0 (CITE-seq).

Each assay technology measures two modalities. The Multiome kit measures chromatin accessibility (DNA) and gene expression (RNA), while the CITE-seq kit measures gene expression (RNA) and surface protein levels.

### Data Pre-Processing

#### Multiome data

##### snRNA

Standard quality control (QC) metrics (total counts, number of detected genes, and percent mitochondrial RNA) were computed, and cells were filtered using thresholds optimized by visual inspection of diagnostic plots. The applied thresholds were: total UMIs between 3,100 and 40,000; at least 2,000 detected genes; and a mitochondrial RNA fraction of 20% or less. Genes expressed in fewer than 20 cells were removed. Counts were normalized per cell to 10^6^ and log-transformed with log1p. Doublets were detected in the RNA modality with Scrublet (run per library,expected doublet rate was set to 10%) via the Scanpy interface^64^. Doublet scores were smoothed over the 15-nearest neighbor graph, and cells with a smoothed doublet score of 0.1 or higher were removed. This threshold was selected after scanning values from 0.1 to 0.4 in increments of 0.02 and visually inspecting results.

##### scATAC

Library quality was assessed via transcription start site (TSS) enrichment and fragment length distributions, after which cell-level filters were applied: total fragment counts between 1,000 and 80,000; at least 750 features per cell; TSS enrichment of at least 0.8; and a nucleosome signal of 2.0 or less. Features present in fewer than 20 cells were removed. Accessibility was binarized, normalized using term frequency–inverse document frequency (TF–IDF) with a scale factor of 10,000, and reduced in dimensionality via truncated singular value decomposition (LSI).

After QC, only barcodes present in both modalities were retained for the final data sets.

### CITE-seq data

#### scRNA

The same QC metrics and plotting workflow were applied as for the Multiome RNA modality. The thresholds used were: total UMIs between 10,000 and 50,000; at least 2,000 detected genes; and a mitochondrial RNA fraction of 20% or less. Genes expressed in fewer than 20 cells were excluded. Doublets were identified in the RNA modality as in the snRNA-seq modality in the Multiome data.

#### Surface proteins

Raw antibody-derived tag (ADT) counts were normalized using DSB-normalization^65^. Non-human isotype controls (mouse/rat IgG) and an empty-droplet log10-UMI range of 1.5–2.8 were used for background estimation. Detection was verified by comparing mean expression of human targets versus non-human controls.

Only cells passing QC in both RNA and protein modalities were retained.

#### Splits

Following the usual structure of machine learning competitions, each data set is split into a training set, a private test set, and a public test set. For the training set, both model inputs and the correct predictions are released - allowing participants to train the model. For the private and public test set, only model inputs are released and participants have to make predictions on these. For the public test, participants can submit predictions prior to the end of the competition, and immediately see the result - allowing competitors to understand the performance of their model and other submitted solutions.

The data splits are arranged as follows: The training set comprises samples only from donors 1-3. The public test set contains samples only from donor 4. The private test set consists of samples from all donors 1-4 but from an unseen day. For the Multiome samples, the training set comprises samples only from days 2, 3, 4, and 7. The public test set contains samples only from days 2, 3, and 7. The private test set comprises data only from day 10. For the CITE-seq samples, the training set and public test set consist of samples from days 2, 3, and 4. The private test set comprises samples only from day 7. There are no day 10 CITE-seq samples in any split.

#### Evaluation metric

The average Pearson’s R between predicted and true values within each cell was used as the evaluation metric. Due to the different number of observations (cells) of the test sets for each subtask, the performance in predicting surface proteins was weighted higher in the final score (about 0.743 to 0.256). To reduce computational resources needed for evaluation, cells were randomly subsampled to 16780 cells.

#### Evaluating robustness of ranking and lower bound optimal performance

To assess the stability of competition rankings, we generated 300 bootstrap samples of the top 50 submissions by sampling predictions with replacement across cells and targets, recalculating the Pearson correlation for each. To estimate a lower bound on optimal predictive performance, we trained a k-nearest neighbors (kNN) regression model using the entire dataset, including the private test set. This model used k=100 and predicted each cell’s output by averaging the target values of its 100 nearest neighbors in PCA space, based on Euclidean distance.

#### Evaluation of validation strategies

We created 12 validation sets for the CITE-seq task using popular or well-performing approaches from the competition. For random validation sets, ⅓, ⅕ and 1/10 of the cells in training data was used, which corresponded to 20051, 12853 and 6749 cells. The same cells were used for validation of all models in our experiments. For testing day-out or donor-out strategies, we put all cells from the furthest time point or from a particular donor to a validation set. Validation set using all cells from the day 2 consisted of 21942 cells, for day 4 there were 28145 cells, sets excluding donors D2m and D3m consisted of 24803 and 23986 cells correspondingly, and combined validation strategies excluding both one day and one donor used 38350, 38452, 42799, and 42620 cells for day-2-D2m, day-2-D3m, day-4-D2m and day-4-D3m correspondingly. Finally, for evaluating adversarial validation strategy, we took the same 5997 cells that rank 3 competitor used in his solution. To obtain this validation set, the competitor trained a LGBM model to predict whether cells come from training or test set. Part of the transcriptomics dataset labeled “X_test_best_128” in the **Supplementary table 6** was used as a model input. The LGBM model with parameters `boosting=’gbdt’, max_depth=7, min_data_in_leaf=20` was trained with learning rate 0.05 to maximize area under ROC curve for classification of training and test data. Then for each donor in training data, 1999 cells with the highest predicted probability of belonging to the test set were selected as the validation set. Almost 80% of adversarial validation set consisted of cells from day 4 followed by cells from day 3 and with almost no cells from day 2. All 3 donors from the training data were evenly represented in this validation set.

We compared 23 variants of CITE-seq models from top 3 competitors using these validation strategies (**Supplementary Figure 3a**). For rank 2 model, which averaged predictions of 2 different neural networks (**Fig. 3c**), we evaluated performance of each of the networks. For rank 1 CITE-seq model, we tested original model with different random seeds (namely, 0, 1, 2, 3, 42, 73, 666, and 1998), 1 and 3 decoder blocks instead of original 5 (see Supplementary for a detailed description of the model), ReLU activation instead of original GELU, 1024 neurons in the fully connected layers of both encoder and decoder instead of 2048 in the original model, and a combination of ablations, consisting of reducing the number of neurons in fully-connected layers to 1024, number of decoding blocks to 3, and using ReLU activation in all layers. For rank 3 CITE-seq solution, which was based on a linear model on top of concatenated outputs of 3 fully connected neural networks with ReLU or Mish activation function and an extensive preprocessing of data, we simplified the model to a single neural network. We used neural networks with 1, 2 or 4 hidden layers with layer normalization and ReLU activations. Hidden layers contained the following number of neurons (labels according to **supplementary figure 3a**): 128 for OneLayerPerceptron; 256 for OneLayerPerceptronV2; 256 and 128 for ThreeLayersPerceptron; 256 for both hidden layers of ThreeLayersPerceptronV2; 256 and 3 layers of 128 neurons for FiveLayersPerceptron; 256, 200, 180, and 128 for FiveLayersPerceptronV2. The final layer of all models was linear with 140 neurons matching the number of surface proteins in the CITE-seq data.

To calculate the quality of validation strategies, every model variant was trained on each training set and evaluated on the validation set. Similarly to a common approach at the competition, models were then retrained on the entire training data and used to predict values for the test sets. We used Spearman correlation coefficient to compare ranking of models by their validation score and by the ground truth private test score (**Fig. 3f, supplementary figure 3a**).

#### Ablation study of top-performing models

We dissected the best approaches per task to understand what makes them successful. For the best Multiome prediction model from rank 1 competitor (**Fig. 3a**), we tested how simplifying the model, removing parts of the input data or ablating pre- and postprocessing steps affects the performance. On the model side, we tested removing MSE loss for predicted residuals; model without 1st or 2nd head; using ReLU activation function instead of GeLU in all layers; using batch normalization or no normalization instead of layer normalization; reducing the number of decoder blocks to 3 or 1; decreasing width of fully-connected layers from 2048 to 1024; removing per-median division of input data or targets, and the combined ablations including removing the 1st head of the model and leaving only 1 decoder; removing 1st head and MSE loss in the 2nd head; removing 1st head, MSE loss in the 2nd head, and leaving only 1 decoder block. We then followed the original training procedure, obtained the prediction of gene expression and evaluated it with Pearson’s correlation to the ground truth on the private test set (**Fig. 3b**), following the structure of the competition. In total, we performed 14 ablation experiments for the winning Multiome prediction model.

For the top performing CITE-seq approach from the rank 2 competitor, we noticed that predictions of 2 underlying neural networks (**Fig. 3c**) have similar performance (**Fig. 3d**). Given a simpler architecture of the model using MSE-loss, we focused our ablation study on this model only. We tested changing Bidirectional GRU block to LSTM or to a simple linear layer or completely removing it; using dropout with fixed parameter or no dropout at all instead of Gaussian dropout; reducing width of fully connected layers to 1000 or to 500; reducing the number of fully connected layers from 3 to 2 or to 1; and changing Swish activation functions to ReLU. As the original solution used 8 differently pre-processed datasets, including LGBM predictions (**Supplementary table 2**), we tested how ablating these datasets changes the prediction quality. We one-by-one removed each of the 4 datasets using predictions of LGBM models; datasets using raw count data; datasets using CLR-transformed expression data (number 1 in the **Supplementary table 2**); raw data-based dataset enriched by selected features (number 4 in the **Supplementary table 2**); all LGBM predictions; and all LGBM and raw counts-based datasets (numbers 3-8 in the **Supplementary table 2**). Ablated models were trained in the same way as the original one. This included random splitting of the training data into 5 folds, training the model on 4 folds, selecting the best model parameters according to validation loss on the validation fold, using it to predict the test data, and averaging prediction of 5 models. Predicted values were then compared with the ground truth on the private test set using Pearson’s correlation. In total, we performed 22 ablation experiments for the top-performing CITE-seq prediction model.

### Evaluating integration of prior biological knowledge

#### Integration of prior biological knowledge in CITE-seq task

To assess the impact of prior biological knowledge, we constructed input features by concatenating data-driven and biology-informed representations of gene expression.

#### Data preprocessing and input features

Raw gene expression matrices were normalized using SCTransform (v0.3.2), and each gene was standardized to zero mean and unit variance. Dimensionality was reduced via truncated singular value decomposition (SVD), using the top 128 left singular vectors multiplied by their corresponding singular values. Gene features were encoded for each of the 134 target proteins, corresponding encoding genes were identified using MyGeneInfo queries. Expression values of these genes were extracted from the normalized matrix and used directly as features. A mapping between protein names and Ensembl gene IDs is provided in **Supplementary Table 7**. Network-propagated features were derived for each encoded gene by propagating over gene interaction networks using random walk with restart (RWR). The propagation followed the recurrence:

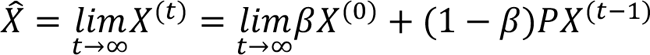

where *X*^(0)^ is the normalized gene expression matrix, β ∈ (0, 1) is the restart parameter that is set to 0.85 in our experiments, and *P* is the column stochastic matrix derived from the gene interaction adjacency matrix. The propagation process is repeated until convergence, resulting in a smoothed gene expression matrix *X^^^*. We evaluated networks from STRING v11 (coexpression, experimental, database, and text-mining evidence) and HumanBase (v2021) (generic and tissue-specific, including bone marrow and HSC-specific networks). Network propagation was performed for 0 to 5 hops, and features from each step were concatenated to the input feature set.

#### Model architecture, training and evaluation

The final input feature set was the concatenation of: (i) 128 SVD components, (ii) encoding gene expression, and (iii) network-propagated features. This was passed to a 2-layer multilayer perceptron (MLP) with 512 hidden units per layer, ReLU activation, and dropout of 0.1. Models were trained using mean squared error (MSE) loss, the Adam optimizer (learning rate = 1e−3), and a batch size of 256 for 100 epochs. We evaluated models using batch-aware cross-validation, where each fold corresponded to a unique donor-day combination. Pearson correlation between predicted and observed protein abundance was computed for each target and averaged across targets and folds. Final performance was assessed on held-out public and private test sets by retraining on the full training data.

#### Integration of prior biological knowledge in Multiome task

To test the utility of databases containing gene regulatory interactions, we combined SVD-derived ATAC features with multiple biologically informed feature sets representing different regulatory mechanisms.

#### Data preprocessing and input features

Chromatin accessibility matrices were TF-IDF normalized and standardized. The top 512 components from truncated SVD were used as baseline features. Accessibility scores for each gene were computed as the sum of peak counts within the gene body and upstream 2 kb region, using the default parameters from Seurat::CreateGeneActivityMatrix (v4.1.1). Only protein-coding genes were considered. To derive enhancer-gene associations, peak counts from annotated enhancer regions were aggregated per gene using enhancer-gene mappings from the following databases: scEnhancer^48^, SEdb^66^, ENdb^67^, SEanalysis^68^, and dbSUPER^69^. Only enhancer-gene pairs relevant to hematopoietic stem cells (HSCs) were retained.

eQTL features were extracted by overlapping peaks with expression quantitative trait loci (eQTLs) from GTEx v8 (whole blood) and mapping them to target genes. Accessibility scores from these eQTL regions were aggregated per gene. Only lead eQTLs with FDR < 0.05 were considered. All biology-informed features were aggregated into gene-level matrices. To ensure fair comparison, only the intersection of genes with valid annotations across all feature types was used (n = 1,115). Feature matrices were concatenated with the corresponding SVD components.

#### Model architecture, training and evaluation

We trained 2-layer MLPs (512 units per layer, ReLU activation, dropout = 0.1) on the combined input. Models were optimized using Adam (learning rate = 1e−3), with MSE loss and batch size of 256 for 100 epochs. Seven-fold cross-validation was performed using a leave-donor or leave-day-out strategy. Final performance was assessed on held-out public and private test sets. We also computed principal component regression (PCR) scores to quantify the amount of biological variance captured by each feature set, using cell type, day, and donor as covariates as implemented by Luecken et al^70^.

#### Computation and analysis of SHAP values

Since SHAP was not initially designed for use with ensemble models, we made the assumption that SHAP feature attributions are additive with respect to the same input. That is, SHAP values computed using the same input in two different models in which the predictions are linearly added, can also be summed to derive the total importance. Briefly, we used SHAP to compute a feature importance for each of the 84 hand-selected features and the 300 SVD features. We then reverse transformed the feature importances of the SVD features to derive feature importances per RNA (**Methods**), and examined the results of the top ranking RNAs.

Makotu’s ensemble contained 18 multi-layer perceptrons, with two models, numbered 16 and 17 in **Supplementary table 6**, outperforming most models in the ensemble, despite simple data preprocessing. Thus, we decided to focus feature attribution experiment on these two models. SHAP was applied to both MLPs using the provided model weights. For background data in SHAP, we used the median RNA expression or the median value of the TruncatedSVD feature. Model 16 incorporated 128 SVD features, while model 17 used 64. As standard, feature attributions were computed on the test set^71^. To streamline computation, given the time-intensive nature of SHAP and to minimize cell type bias, we subsampled each cell type to 160 cells and trained KernelExplainer accordingly. SHAP values for each pair of input and output features for each data point in the test set were generated using KernelExplainer().shap_values, with the nsamples parameter set to 300. We then used the inverse_transform() function of TruncatedSVD to reverse the transformation of the SVD features to convert attributions from TruncatedSVD space back to the original 22,001 RNA expression space. The attributions of the hand-selected 84 genes were not modified. Final attributions values for all genes were taken by computing the mean attribution value between model 16 and 17. We deemed this to be an acceptable method of ensembling attribution scores between ensemble models as 94% of attributions between the two models had the same directionality (e.g. increasing the RNA expression value increased the attribution score in both models). Finally, genes were ranked by their mean attribution values across all test data points, as recommended by SHAP (beeswarm plot), for a single protein target at a time.

Gene set enrichment was run using GO:BP^72^ and the gprofiler-official python package^73^, with all RNAs used as background.

#### Donor label mapping

For better readability, in the publication we changed the donor labels. They correspond to the labels in the data as follows: D1f – Donor 13176, D2m – Donor 31800, D3m – Donor 32606, D4m – Donor 27678 (test).

## Data availability

Processed data for this study is available on GEO at accession GSE305370. The corresponding FASTQ files are available on SRA under BioProject PRJNA1300406.

## Code availability

The code is available at https://github.com/lueckenlab/OpenProblems2022Analysis. Reimplemented O2C1 model and preprocessing is available at https://github.com/lueckenlab/senkin-tmp-cite-pred.

## Acknowledgements

We thank all competition participants for their valuable contributions and discussions, and the Kaggle platform for hosting the competition. We acknowledge Allegra Lord, Aakash Kabbin, and Sakina Saif for their assistance with data processing for the competition. We are grateful to Nikita Moshkov for providing feedback on an earlier version of the manuscript. We further acknowledge the support of Cellarity and Data Intuitive, whose contributions facilitated the data generation, competition and benchmark set up. V.A.S. and C.L. are supported by the Helmholtz Association under the joint research school “Munich School for Data Science - MUDS”. This project has been made possible in part by the Chan Zuckerberg Initiative DAF, an advised fund of Silicon Valley Community Foundation.

## Author contributions

C.L., V.A.S., and M.D.L. conceived the study. P.H., R.C. and D.B. organized the competition and P.H., B.D., and M.C. generated and prepared the data for the competition. C.L, V.A.S., H.W., Y.J., P.H., R.L., W.T., and Y.W performed data analysis and implemented, together with W.T., K.W., and R.C., computational methods. Y.X., J.T., and D.K. contributed to supervision and provided resources. M.L., D.B., and F.T. supervised the project, provided funding, and contributed to overall study design and administration. C.L., V.A.S., M.L., H.W., and P.H. drafted the original manuscript. All authors reviewed and approved the final manuscript.

