## Supplementary Information for "Longitudinal modality prediction learns gene regulatory patterns: insights from a single-cell competition"

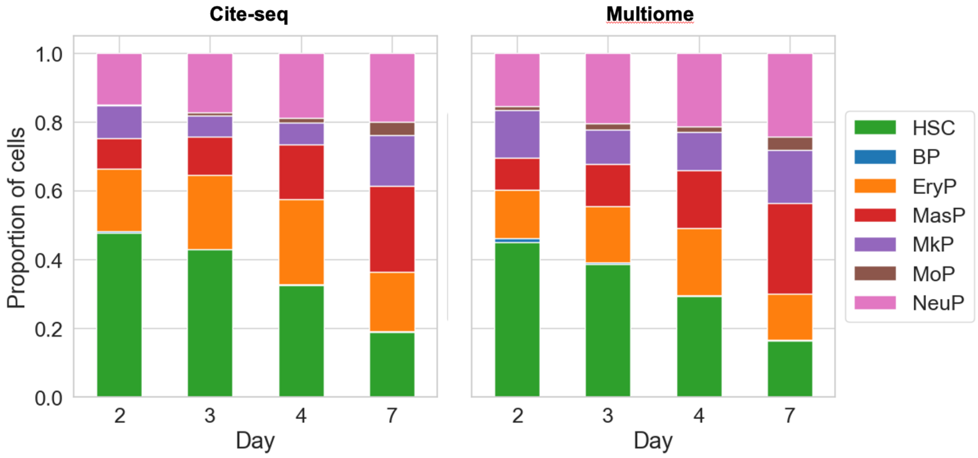

**Supplementary figure 1.** Cell type composition over time. This shows a reduction of hematopoietic stem cells (HSC) while the amount of B-Cell progenitors (BP), erythrocyte progenitors (EryP), mast cell progenitors (MasP), megakaryocyte progenitors (MkP), monocyte progenitors (MoP) and neutrophil progenitors (NeuP) increases.

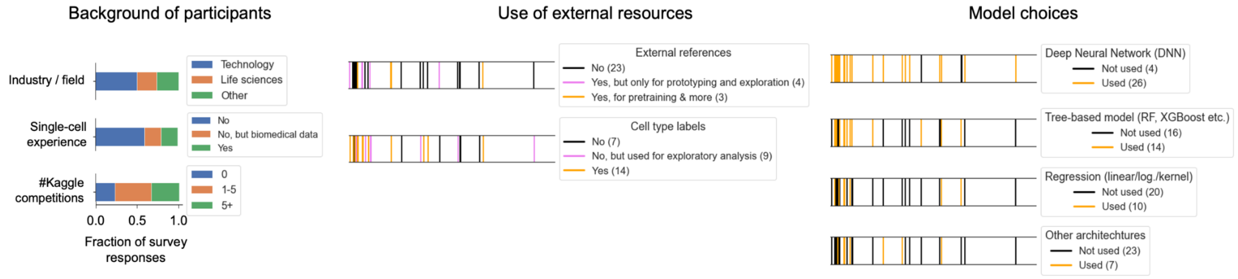

**Supplementary figure 2. Survey results.** Shown are relevant results of a survey, participants were asked to fill out after the competition was finished.

#### Top 1 solution: neural network is all you need

The winning submission from a participant called Shuji Suzuki achieved superior performance in solving the Multiome task (Figure 2d), and ranked 2nd in the CITE-seq task (Figure 2e). It holded the place 27 on the public leaderboard at the end of the competition and rose to the top after the evaluation on the private test set. This solution is based on a comparingly simple NN-based model, which, as we show later with ablation study, can be simplified even more (Figure 3c,d).

A distinctive feature of this solution is the extensive preprocessing of the data (**Supplementary Table 1**), both for the models' input and for the prediction targets. The dimensionality of Multome task was too high to work with the full data so the common approach among many competitors was to reduce its dimensionality with Truncated Singular Value Decomposition (TSVD) both for chromatin accessibility and RNA data. For Multiome task, the 1st place solution used preprocessed and TSVD-transformed chromatin accessibility data as well as aggregated cell- and sample-level statistics (**Supplementary Table 1**) as an input for a multilayer perceptron. 2 blocks of linear layer followed by GELU[74] activation function, dropout and layer normalization, denoted by the author as an "encoder", preceded a sequence of "decoder" blocks each containing 2 modules, described here as "heads" (Figure 3c). The first head uses linear layer to predict 128 TSVD-values of the target RNA expression, which are then converted to counts using a reverse transformation. The second head predicts RNA-counts residuals for every gene of 23,418 present in the dataset. This prediction is added to the output of the first head, thus correcting it. The first mechanism ensuring this is the correlation loss between the ground truth counts and the sum of inversed predicted TSVD values and residuals. Additionally, MSE loss is used to force residuals to be similar to the difference between target counts and inversed predicted TSVD values. The sum of predictions from 2 heads, i.e. the corrected prediction, is used as an input for the next decoder block with identical architecture. The final result is the aggregation of the predicted RNA counts matrix from each of 5 decoder blocks.

Interestingly, we found that this architecture can be substantially simplified while still keeping great performance. When ablating individual parts of the model or preprocessing steps, we noticed that most of them were redundant (Figure 3d). In particular, normalization layers can be changed to the batch normalization (score on private test set for Multiome $P_m$ = 0.5781) or completely dropped ($P_m$ = 0.5765), activation functions changed to a simpler ReLU ($P_m$ = 0.5779), number of decoder blocks decreased to 3 ($P_m$ = 0.5769) or even 1 ($P_m$ = 0.563), or MSE loss for residuals prediction can be cut ($P_m$ = 0.5784) without significant decrease of the performance comparing to the original model ($P_m$ = 0.5782). Even the combination of ablations, namely using only 1 correlation loss for the corrected prediction instead or original 3, and having only 1 decoder block instead of original 5 resulted in the model, which would still put the solution on the first place ($P_m$ = 0.5749). Removing the per-cell median division in the prerocessing step did not impact the performance neither for input ($P_m$ = 0.5770) nor for targets ($P_m$ = 0.5770). The only ablation that significantly decreased the solution quality was removing the second head in the decoder block ($P_m$ = 0.5388), suggesting that prediction of the full genes set was a key for the model performance. It is worth highlighting that while the original submission included ensembling of the models trained with different random seeds and fine-tuned on the different batches, in the ablation studies we only used a single model but the performance was still on par with the original solution.

The CITE-seq approach for the 1st place solution is quite similar in terms of extensive preprocessing, metadata usage, and modeling. It is worth noting that in addition to using TSVD-transformed RNA counts, which was typical for many competitors, important features selection was leveraged. Both data-driven approach with selecting highly correlated features and prior knowledge from Reactome Pathway database was used (see **Supplementary Table 1** for details). The selected genes values were concatenated together with TSVD-transformed RNA-seq dataset and metadata to use as a model input. Interestingly, despite comparably low number of surface proteins (140), which allowed direct prediction of every feature, the target data dimensionality in this solution was also reduced to 128 components with TSVD.

Potentially, it allowed the model to easier predict abundance of the correlated proteins. The model architecture is almost identical to the model for Multiome task but does not contain residuals prediction. Instead, the output of the first head is compared with TSVD-transformed targets using MAE loss, and the second head predicts every protein level, which are then subject to the transformation reversing preprocessing and compared with ground truth using the correlation loss.

**Supplementary Table 1. Preprocessing of the model input and targets for rank 1 solution. Inputs are chromatin accessibility values in a region and RNA counts, while targets are RNA counts and surface protein abundance levels for Multiome and CITE-seq tasks respectively. Metadata refers to additional information about donors and cells as well as to aggregated statistics**

| Task | Input preprocessing | Target preprocessing | Metadata usage |
| --- | --- | --- | --- |
| Multiome | 1. Transform the data with TF-IDF<br>2. Divide TF-IDF values by non-zero median values per cell<br>3. Reduce the dimensionality with 128 components truncated SVD | 1. Divide raw counts by non-zero median value per cell<br>2. Perform log1p transformation<br>3. Perform TSVD transformation and reverse transform, only take restored values where original data had 0 counts to impute them<br>4. Transform values to Z-scores per cell<br>5. Subtract median values per-feature<br>6. TSVD-transform the data | The following statistics were additionally used as an input for the models<br>● 1, 2, and 3rd quartiles of the non-zero values per cell<br>● Mean and standard deviation of cell features<br>● Cell type counts and percentages per donor and timepoint<br>● Ratio of regions with non-zero accessibility per chromosome and same ratios log1p-transformed<br>● Per-column z-scores of the above-mentioned features<br>● Sex of a donor |
| CITE-seq | 1. Divide counts by non-zero median values per cell<br>a. Select genes that have a significant Spearman correlation (abolute value of a coefficient $\geq$ 0.1, p-value $<$ 0.01) in at | 1. Transform values to Z-scores per cell<br>2. Subtract median values per protein<br>3. Reduce dimensionality with 128 components | Same as for Multiome except for per-chromosome ratio of non-zero values |

| | | |
|---|---|---|
| | least 8 donor-timepoint pairs from 12 available in the train set. There were 71 such genes<br>b. Select genes related to target protein using a Reactome pathway database. 3 genes for each target protein with the smallest median p-value of a correlation test were selected among the genes with abolute value of a Pearson correlation coefficient ≥ 0.2, p-value < 0.001 in at least 8 donor-timepoint pairs<br>c.<br>i. Impute 0 counts with 128 components TSVD, analogously to Multiome<br>ii. Subtract median values per gene<br>iii. Reduce dimensionality with 128 components TSVD | TSVD |

### Top 2 solution: data diversity allows model simplicity

The second-place solution was presented by two participants nicknamed senkin13 and tmp (baosenguo). It is notable for its outstanding CITE-seq task performance (Figure 2c) and robustness, confirmed by the 1st place on a public leaderboard and 2nd place on the private test set evaluation. Commonalities between CITE-seq and Multiome models include training neural networks on variously preprocessed datasets and on the output predictions of simpler machine learning models. While it was on par with other top 50 solutions for the Multiome task (Figure 2d), this approach performed outstandingly well for the CITE-seq task (Figure 2e).

For Multiome prediction, TF-IDF preprocessed counts were transformed with 100 components TSVD and then converted to Z-scores per cell. These values were used as an input to LGBM model predicting 1000 TSVD components of RNA data. Values predicted with LGBM were once again TSVD transformed to 100 components and used as a dataset for subsequent models. Additionally, raw chromatin accessibility counts were transformed using Centered Log-Ratio (CLR)[75], then compressed 100 components TSVD and converted to Z-scores cell-wise. 2 neural networks were used to predict all 23418 genes values using CLR-transformed values and compressed LGBM predictions as an input. Both networks contained 3 fully connected layers (600 and 500 neurons in each layer for networks 1 and 2 correspondingly) with Swish activation function and Gaussian dropout with parameter 0.3 in between. The first NN used cosine similarity loss, while the second was trained with Huber loss with the parameter delta = 0.4 to predict per-cell Z-scores of the genes' RNA expression. The predictions of both networks were averaged with equal weights. This approach led to place 23 in the Multiome task. To note, in the solution description the authors claimed other machine learning models to be used as well, but we did not find the confirmations in their code.

For CITE-seq prediction, the team members produced versions of the dataset using diverse preprocessing (**Supplementary Table 2**). TSVD-transformed raw and normalized counts, raw values of genes selected by high correlation with target and of genes encoding target proteins, as well as predictions of 4 LGBM models using various datasets and hyperparameters, were used as an input to neural network. Two similar deep learning models were then trained to make the final predictions, one using MSE-loss and another using cosine loss further referred to as MSE-model and cos-model (Figure 3e). Both architectures had a Gated Recurrent Unit (GRU)[76] either as the first layer in the MSE-model or as the last layer prior to the output linear layer in cos-model. The core of the networks contained fully connected layers followed by either swish or exponential linear unit (ELU)[77] activation and Gaussian dropout with average drop probability 0.1 and 0.2 in MSE- and cos-models, respectively. MSE-model simply contained sequential layers, while in cos-models, outputs of each linear block were not only passed further but also concatenated to the input of the last linear layer. Outputs of two networks were averaged with weights 0.45 and 0.55 to make the final prediction.
To explain the success of this model in CITE-seq data prediction, we performed a series of ablations. First, we completely removed one of two models and obtained almost identical scores – 0.8467 for the cosine loss model and 0.8463 for the MSE loss model compared to 0.8477 for the original model. We then focused our ablations only on the MSE-model due to its lower complexity. Most of the architecture ablations did not significantly decrease the score, including changing GRU to another recurrent Long Short Term Memory (LSTM) block, to a linear layer or even completely removing it; as well as changing dropout to standard instead of Gaussian or dropping it completely, and reducing the number of FC layers to 2 or 1, or changing their width to 1000 or 500 (**Fig. 3d**). Replacing swish activation with ReLU resulted in a worse prediction (score on private test set for CITE-seq $P_c$ = 0.8452) suggesting its importance for this architecture. Removing most individual datasets from the input did not substantially worsen the results, however, removal of dataset #8 containing LGBM predictions for unnormalized protein levels from the normalized expression of all genes resulted in a worse score not outbeating the other top participants. When all datasets using LGBM predictions were ablated, the score reduced to 0.8447, putting the prediction on place 10 for CITE-seq and 10 in the overall ranking. If datasets using raw data were dropped in addition to LGBM predictions, the score fell down to 0.8436, making this model on par with many other CITE-seq prediction approaches (**Fig. 2e**). These results suggest that the data preprocessing diversity was the key to explaining this model performance and its architecture could be substantially simplified while still keeping the great score.

**Supplementary Table 2**. Preprocessing strategies used by the best CITE-seq prediction model

| Dataset number | Preprocessing steps | Number of features |
|---|---|---|
| 1 | 1. CLR-transformation of the raw RNA expression data<br>2. 200 components TSVD transformation | 200 |
| 2 | 1. Division of raw counts data by average expression per cell<br>2. Taking a square root of the data<br>3. Conversion to Z-scores per gene<br>4. Batch effect correction by subtracting the median value of expression of each gene on a given day of differentiation<br>5. 100 components TSVD transformation | 100 |
| 3 | 64 components PCA transformation of the dataset #2 (prior to TSVD transformation) | 64 |
| 4 | 1. Convert target values to Z-scores per cell<br>2. Using raw RNA data, select 10 genes most correlated with each target<br>3. Combine selected genes with 144 genes encoding target proteins<br>4. Save raw RNA counts of the selected features | 245 |
| 5-8 | Train LGBM models with varying inputs and hyperparameters to predict abundance (normalized for 5-7, raw for 8) for 140 proteins:<br>5. Used normalized expression values of all genes as an input. Hyperparameters `feature_fraction=0.05`, `bagging_fraction=0.9`, and no regularization (`lambda_l1`, `lambda_l2`) were used.<br>6. Used all datasets 1-4 as input. Hyperparameters `feature_fraction=0.7`, `bagging_fraction=0.7`, `lambda_l1=0.1`, and `lambda_l2=1` were used.<br>7. Used raw expression values of all genes as an input to predict unnormalized protein levels. Hyperparameters `feature_fraction=0.08`, and no regularization (`lambda_l1`, `lambda_l2`) were used.<br>8. Used unnormalized expression values of all genes to predict unnormalized protein levels. Hyperparameters feature_fraction=0.1, bagging_fraction=0.9, lambda_l1=1, and lambda_l2=10 were used.<br><br>Each model prediction was then transformed independently by 100 components TSVD | 4 * 100 |

### Top 3 solution: there is no such thing as not enough ensembling

The participant with the nickname Makotu created the 4th best model for Multiome task prediction and 13th best for CITE-seq, achieving the 3rd place after averaging. His solution was notably robust, holding 11th place on the public leaderboard. We believe that the key feature of this success was the adversarial validation strategy (see Figure 3b and the section on validation) and a wide variety of preprocessing methods and models for prediction. The margins of this paper are too small to fit the description of all the details as for CITE-seq only, 20 models were trained using different combinations of 27 variously preprocessed datasets, aggregating the predictions with different weights obtained by a cross-validation score. An interested reader can find details in the **supplementary table 6** and Makotu's description of his solution[78]. Briefly, for the Multiome task, 26 neural networks with RMSE loss were trained to predict either all genes' expression or 128 components of SVD-transformed RNA data. As an input for the models, combinations of different datasets were used, including 64 components of Latent Semantic Indexing (LSI) of ATAC data, 16 components of SVD-transformed binarized ATAC data, 16 dimensions of word2vec transformation of top 100 regions with the highest counts, 16 dimensions of SVD-transformed features aggregated per 23 leiden[79] clusters, and 16 components of SVD transformation of adjacency matrix between cells. For models predicting the SVD components of RNA data, the inverse transformation was applied before producing the final prediction through weighted averaging.

For CITE-seq prediction, various preprocessing of RNA expression data and metadata were used. In addition to using normalized and SVD-transformed data, the competitor prepared several datasets using word2vec transformation, cell type ratios for each donor, data aggregated per cell type and per clusters obtained with spectral clustering, and features of the similar cells weighted by cell similarity. Notably, two sets of important features containing 84 and 36 genes most correlated with target proteins were used separately without including them in dimensionally reduced data. 18 neural network models and 2 catboost models were trained on different combinations of the datasets. Deep learning models had identical architecture based on 4-layer MLPs containing 3 blocks and a head element. Each block had a linear layer (128, 32, and 8 output features, respectively) followed by layer normalization and ReLU activation function. The output of each block was proceeded to the next block and additionally concatenated to the input of the head, which was used to obtain the final prediction for the values of 140 surface proteins. To understand which models and data preprocessing strategies contributed to the final prediction, we evaluated each model separately (**Supplementary figure 3**). Surprisingly, training the original model only on one of the simplest datasets in the ensemble allowed it to achieve the second best performance among other models trained on a single data source. It contained normalized and log-transformed values of 84 selected genes as well as 128 SVD components of normalized and log-transformed expression data excluding these 84 genes. While this model only scores 0.7721 and would put the competitor at the place 8, it still predicts surface protein information reasonably well, and is drastically simpler than the original solution or any other model we discussed above.

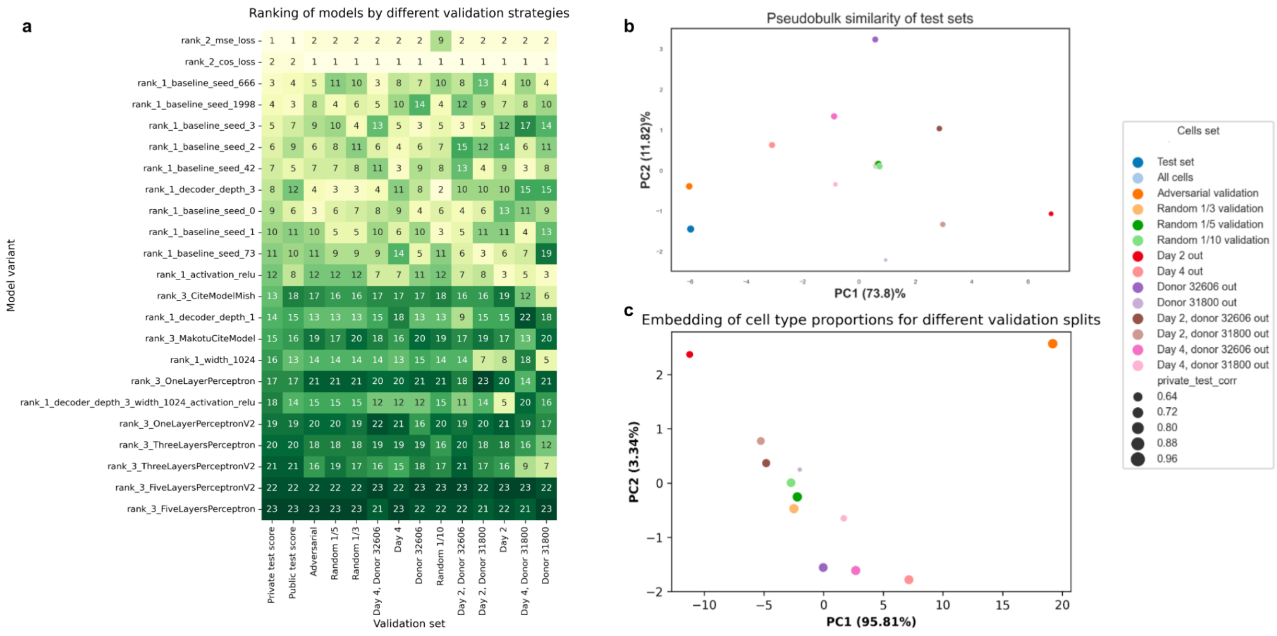

**Supplementary Figure 3.** Validation strategies performance evaluation and interpretation. **a,** Ranking of the CITE-seq model variants of the top 3 competitors by different validation strategies. Models were retrained on subsets of the training data without validation set, and tested on the corresponding validation set. Numbers correspond to the ranking of the Pearson correlation scores between predicted and true protein surface expression for each validation set. Colors represent the ranking of models retrained on the whole training data and evaluated on the private test set. **b,** Transcriptional similarity of validation sets to the private test. Each point is a centroid in TSVD space. Point size represents the value of the Spearman correlation coefficient between the validation score and private test score for different models (Fig. 2b, data from panel a of the current figure). For all cells and the test set, the size is set to 1. **c,** Compositional similarity of validation sets. First 2 components of PCA for cell type proportions are shown. Test set not shown as the labels were hidden.

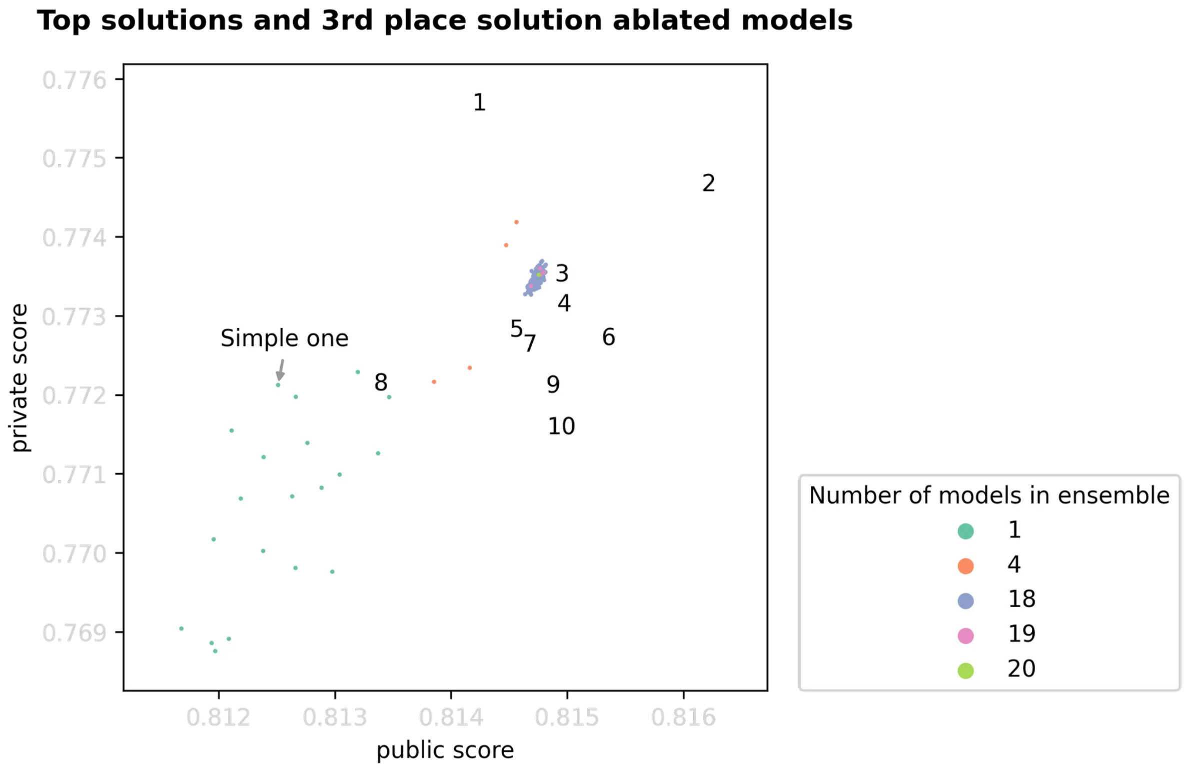

**Supplementary Figure 4**. Distilling successful CITE-seq models from the ensemble of 20 models submitted by the rank 3 competitor. Each point is a prediction by 1 or several models, indicated by color, combined with the same weights as in the original submission. Numbers show the performance of the top 10 solutions of the competition. A simple model marked with an arrow only uses selected 84 genes and 128 SVD components of the rest gene expression data, and yet scores as top 9 in the competition.

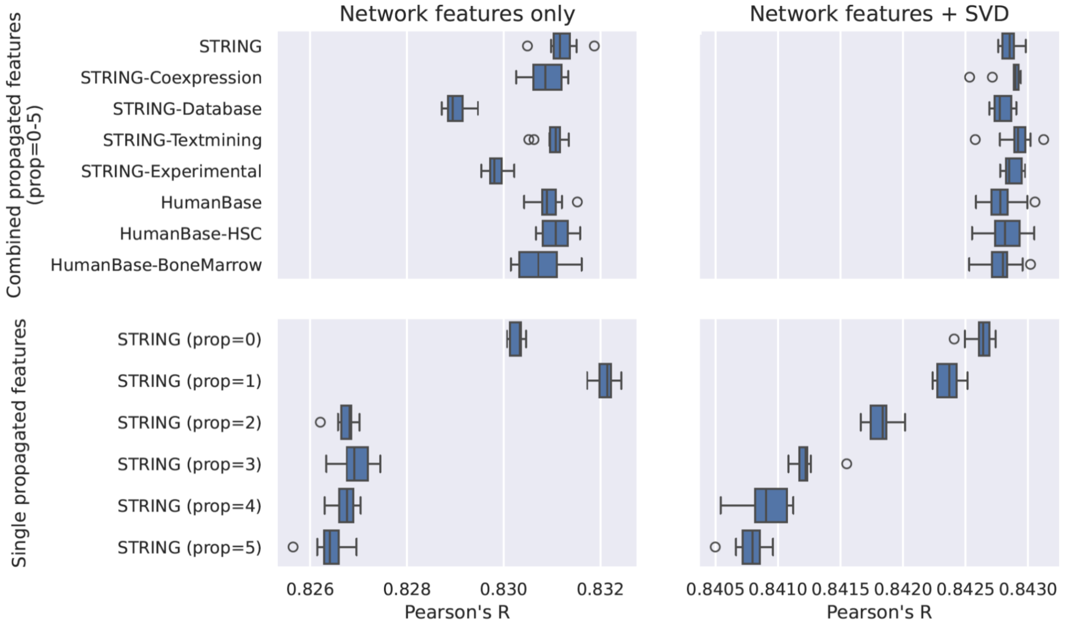

**Supplementary figure 5**. Integration of prior biological information in the CITE-seq task. Boxplots showing the performance of integrating PPI network derived features to predict surface proteins in a simplified 2-layer MLP. To the left prediction performance only using

Network features is shown, on the right these features were combined with SVD components. The top row shows the effect of using different PPI resources, while the bottom row displays the depth of the network propagation considered to derive features for each target protein.

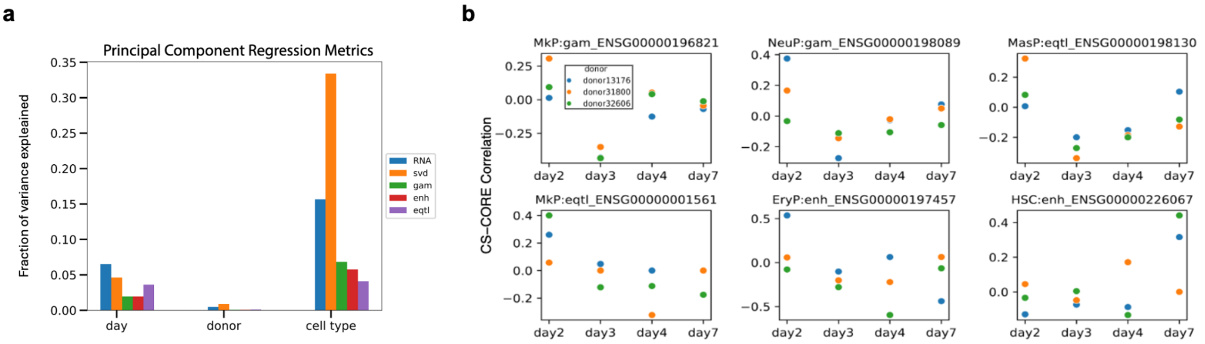

**Supplementary figure 6.** Integration of prior biological information in the Multiome task. **a**, Barplot showing the fraction of variance in the biologically defined features that is associated with time (day), donor or cell type. **b**, Plots showing the correlation of regulatory features with their corresponding target gene expression over time.

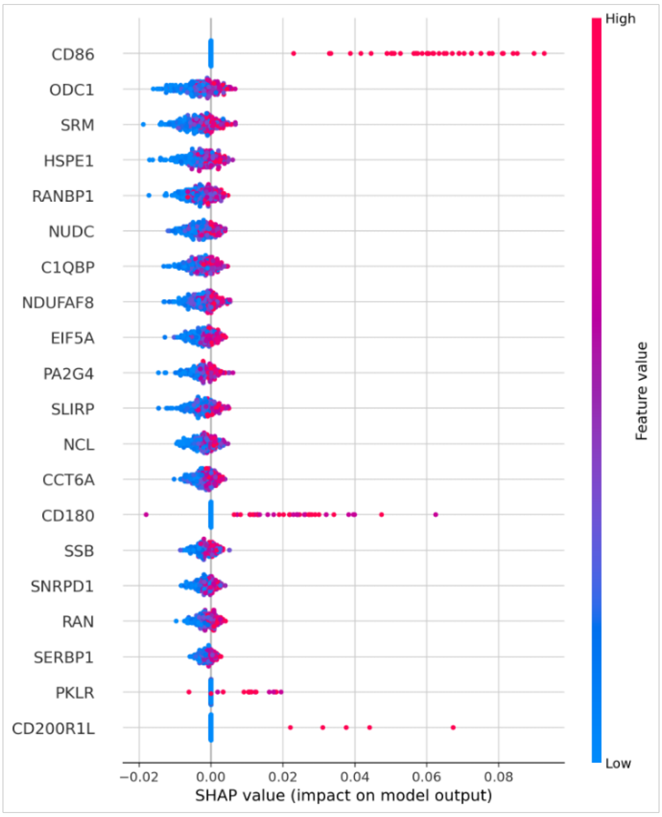

**Supplementary figure 7**. Distribution of average SHAP attribution values aggregated from models #16 and #17 of rank 3 CITE-seq solution. The rows show top 20 most predictive genes for CD86 surface expression. X axis shows feature importance measured as average SHAP value for the two models used. Each point represents one cell.

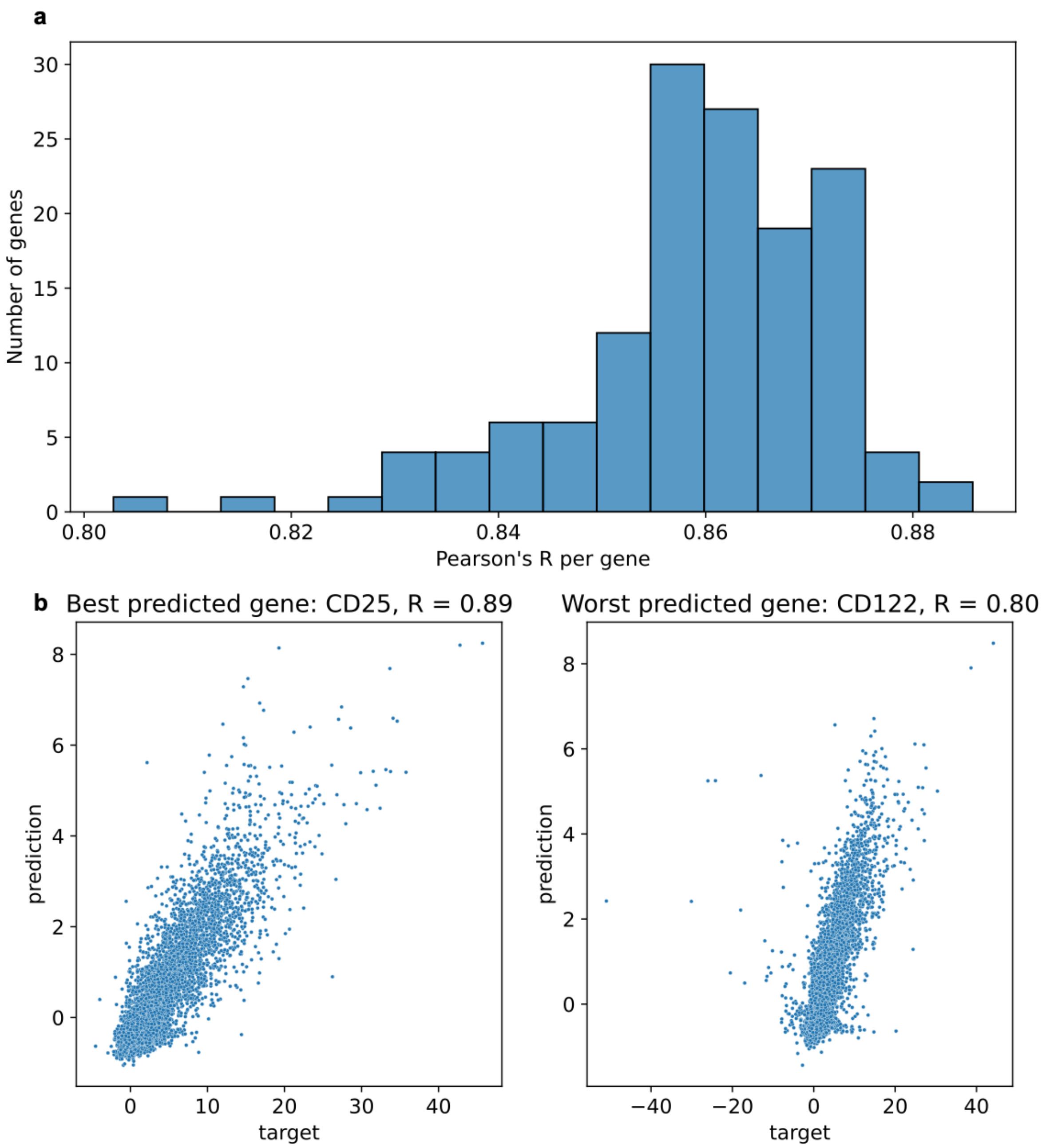

**Supplementary figure 8**. Per-protein assessment of the best CITE-seq model predictions. **a,** distribution of correlation scores per surface protein shows good prediction quality for all proteins in our data. **b,** visualization of predictions for a protein with the highest score (CD25) and for a protein with the lowest score (CD122). Each point represents one cell in the private test set. The title shows Pearson's R score. We can see that the prediction is overall good for both proteins with outliers lowering the score for CD122.
